# Decoding the Valence of Developmental Social Behavior: Dopamine Governs Social Motivation Deficits in Autism

**DOI:** 10.1101/2025.11.12.687985

**Authors:** Xinfeng Chen, Xianming Tao, Zhenchao Zhong, Yuanqing Zhang, Yixuan Li, Ye Ouyang, Zhaoyi Ding, Min An, Miao Wang, Ying Li

## Abstract

The social motivation theory posits that core social deficits in autism spectrum disorder (ASD) arise from impaired social valence assignment during the social critical period, yet the specific dopaminergic mechanisms governing this process remain unclear. We combined high-resolution behavioral sequencing (Social-seq) with fiber photometry to resolve nucleus accumbens (NAc) dopamine during naturalistic juvenile interactions. Sex-divergent social strategies emerged: males exhibited peer play-dominant interactions with action-contingent dopamine release, while females favored environmental exploration with attenuated social dopamine. *Shank3*-deficient juveniles exhibited a triad of dopaminergic dysregulation—blunted signaling during social investigation, pathological inversion during active play, and hyper-responsive to non-social stimuli—recapitulating ASD-like phenotypes. Closed-loop activation of dopamine during play rescued social deficits, establishing a causal link between phasic dopaminergic signaling and social motivation. These findings identify NAc dopamine as a dynamic encoder of social valence and suggest that temporally precise modulation of dopaminergic circuits may offer therapeutic leverage for ASD-related social impairments.

## Introduction

Autism Spectrum Disorder (ASD) is a neurodevelopmental disorder characterized by persistent deficits in social communication and restricted, repetitive behaviors (Vahia, 2013). While early theories emphasized impairments in theory of mind (“mindblindness”) (Baron-Cohen, 1997), growing evidence implicates disrupted social motivation as a core pathological mechanism. The social motivation hypothesis proposes that blunted sensitivity to social rewards in ASD leads to cascading deficits in social orienting, engagement, and learning (Chevallier et al., 2012). These impairments emerge most prominently during juvenile development—a critical period when experience-dependent neural plasticity shapes lifelong social circuits (Foulkes and Blakemore, 2016; Nardou et al., 2019; Towner et al., 2023). However, the neural substrates underlying this developmental vulnerability, particularly the role of dopamine (DA), remain poorly understood.

The mesolimbic DA system, particularly ventral tegmental area (VTA) projections to the nucleus accumbens (NAc), plays a critical role in assigning motivational value to social stimuli (Dai et al., 2022; Solié et al., 2022; Willmore et al., 2022). During typical development, this circuit reinforces prosocial behavior by dynamically encoding the rewarding properties of social cues (Bottini, 2018). In ASD, however, NAc hypoactivity (Shafritz et al. 2015) appears to disrupt this valence assignment process, rendering social stimuli less intrinsically rewarding. This impairment establishes a pathological cycle: diminished social reward sensitivity leads to reduced social attention, thereby limiting opportunities for social learning and further exacerbating social-cognitive deficits (Damiano et al., 2015).

While reinforcement-based interventions like applied behavioral analysis (ABA) have shown clinical efficacy (Dawson and Bernier, 2007; Koegel et al., 2001; Neuhaus et al., 2010), their impact is constrained by three critical neurobiological gaps. First, the precise mapping between developmentally salient social behaviors and their underlying valence remains undefined. Second, the neurobiological basis of the well-documented male predominance in ASD remains unclear (Loomes et al., 2017). Third, it remains unknown whether circuit-based interventions could effectively rescue social motivation deficits during critical developmental periods.

These questions naturally converge on juvenile peer interactions—an evolutionarily conserved behavioral context essential for social circuit development (Del Giudice et al., 2009; Pellis et al., 2010). Rodent studies demonstrate that social deprivation during this social critical period induces lasting socio-emotional impairments (Siviy, 2016; Pellis et al., 2023), mirroring deficits in children with ASD who have limited peer interaction experience (Jordan, 2003). In ASD, disrupted social motivation may initiate a pathological cascade: reduced peer engagement limits crucial developmental social experiences, which in turn exacerbates socio-emotional dysfunction.

This understanding suggests that targeted valence modulation during peer interactions could break this cycle. However, a critical challenge remains: juvenile peer interactions comprise distinct behavioral components, each potentially mediated by unique valence coding mechanisms. Identifying which specific components represent optimal intervention targets remains a critical unanswered question.

To address this, we developed Social Behavior Sequencing (Social-Seq)—an integrated pipeline combining occlusion-resistant 3D pose estimation, self-supervised learning and active learning to decompose naturalistic social behavior into sub-second behavioral “syllables” while simultaneously monitoring NAc core DA dynamics. Applying this approach to wild-type and *Shank3*-deficient juvenile rats (an ASD model) (Meng et al., 2022), we identified sexually dimorphic social interaction strategies and corresponding NAc dopamine responses. Specifically, *Shank3*-deficient juveniles exhibited blunted dopamine responses during social play but heightened responses during non-social exploration. Most notably, closed-loop DA activation precisely timed to play initiation robustly rescued social deficits in *Shank3*-deficient males. These findings demonstrate that NAc-DA dynamically encodes sex-specific social motivational values during development, and that temporally precise dopaminergic interventions can restore typical social function in ASD models.

## Results

### Pose estimation during naturalistic juvenile social interactions

Accurate tracking of social interactions relies on resolving precise body kinematics, a challenging task when juveniles become entangled during play. Traditional multi-animal pose estimation methods (Marshall et al., 2021; Lauer et al., 2022; Pereira et al., 2022; Marks and Mehmet Fatih Yanik 2022 NMI; Han et al., 2024; Klibaite et al., 2025), which attempt direct keypoint estimation from raw video, often fail as overlapping bodies obscure anatomical features. To overcome this, we developed a multi-view segmentation-first strategy implemented through two technological advances.

First, we engineered a nine-camera volumetric imaging system (60 cm arena, Videos S1 and S2) that captures social interactions from multiple angles, thereby reducing keypoint loss due to occlusion (Figures 1A and S1A-S1E; STAR Methods). Second, we implemented instance segmentation to disentangle interacting individuals by training models on rats with distinct fur patterns (Figure 1B). This approach first isolated individual contours before predicting keypoints within each mask, effectively eliminating cross-animal interference during contact-rich behaviors such as pouncing (Figure 1C).

**Figure 1.**
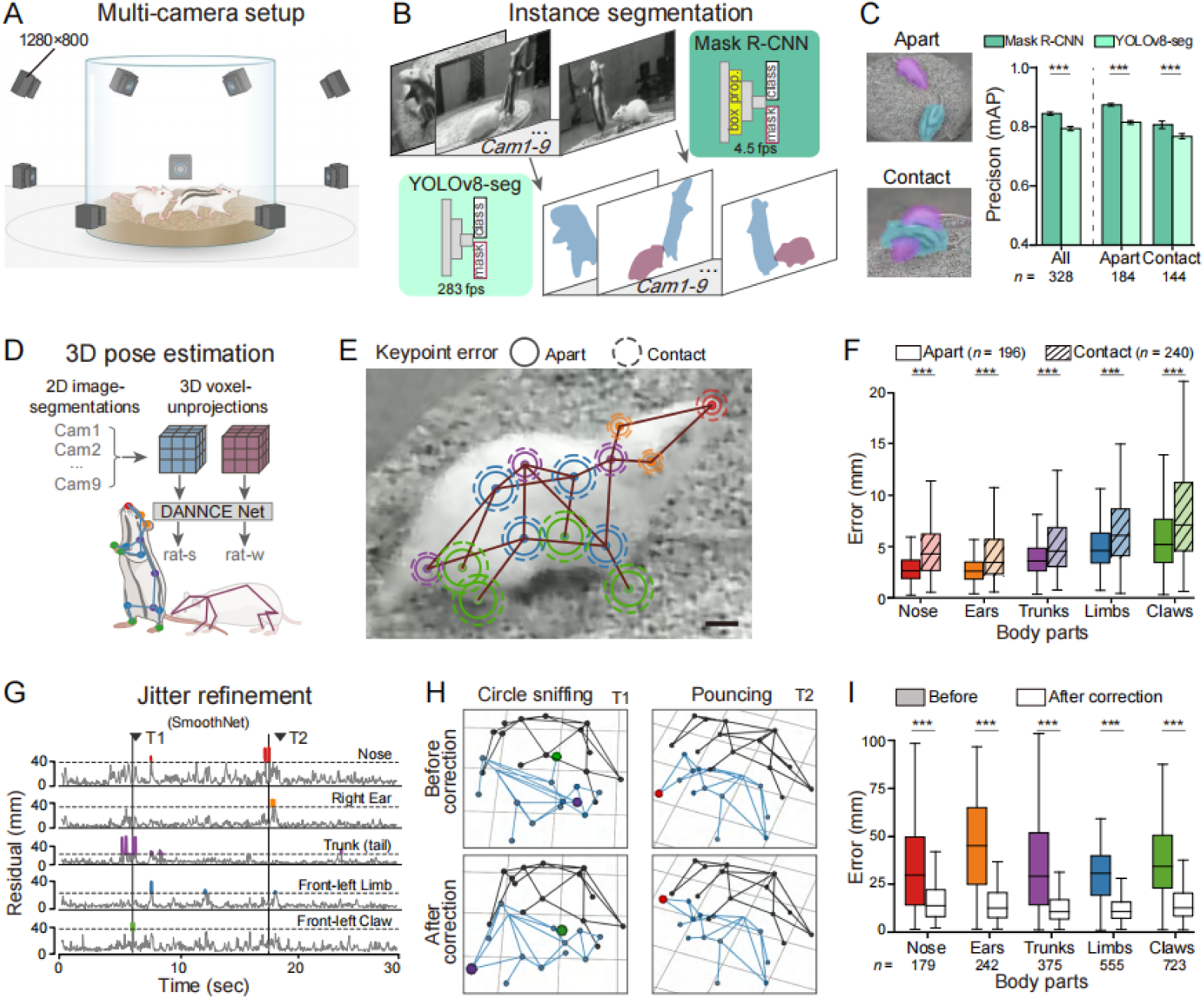
High-resolution 3D pose tracking in freely interacting juvenile rats. (A) Experimental setup. A nine-camera array (synchronized webcams; each at 1280×800 pixels and 120 fps) recorded dyadic social interactions in a cylindrical arena containing corncob bedding. For individual identification, one rat received fur stripe markings (B) Instance segmentation comparison. Bottom left: YOLOv8-seg enables faster processing suitable for real-time applications. Top right: Mask R-CNN achieves more accurate masks via additional bounding-box proposals. (C) Segmentation performance. (Left) Representative contours. (Right) Accuracy quantification for separated (>5 cm apart) *vs.* contacting rats (n = 328 images; paired two-sided t test, mean ± SEM). (D) 3D reconstruction pipeline. Multi-view segmented images were integrated into a voxel volume using DANNCE to track 14 anatomically defined landmarks: nose, ears, trunk (neck/back/tail base), limbs (front/hind), and claws (front/hind). (E and F) Pose estimation accuracy. (E) Spatial error distribution showing median errors (solid circles: separated rats; dashed circles: contacting pairs). Scale bar: 10 mm. (F) Euclidean errors of Mask R-CNN-and DANNCE-derived landmarks relative to manual annotations (two-sided unpaired t test). (G) Jitter detection. Residuals demonstrate deviations between raw and SmoothNet-predicted trajectories. Colored traces (above threshold: > 2-3 SD) indicate jitter events. Triangles mark example timepoints (T1, T2) analyzed in (H). (H) Representative pose corrections. (Top) Original jitter-corrupted skeleton. (Bottom) SmoothNet-corrected outputs. (I) Error reduction following SmoothNet correction for detected jitters (\**p* < 0.05, \*\**p* < 0.01, \*\*\**p* < 0.001; two-sided paired t test relative to manual annotations). See also Figure S1.

Choosing a segmentation approach requires a careful balance between speed and precision. While Mask R-CNN, a region-based convolutional network (He et al., 2017), achieved 84% mask overlap with manual annotations, its processing speed of 4 fps proved insufficient for real-time applications. In contrast, YOLOv8, a single-shot detector (Yaseen, 2024), prioritized speed (283 fps) with a moderate reduction in accuracy (79% overlap), making it ideal for closed-loop experiments (Figure 1B). This modular design allows our framework to adapt to different experimental needs—favoring high precision for offline analysis and speed for real-time interventions.

For pose estimation, we applied DANNCE (Dunn et al., 2021) to track 14 body keypoints from the segmented animals (Figure 1D). The model demonstrated high precision for isolated rats but showed degraded performance during close contact, particularly for occluded body parts such as claws and limbs (Figures 1E and 1F). These errors manifested as biomechanically implausible trajectories with abrupt frame-to-frame keypoint displacements. To address this, we applied SmoothNet (Zeng et al., 2022), a temporal refinement algorithm that integrates multi-frame kinematic relationships to detect and remove motion jitters (Figures 1G-1H and S1F-S1G). The corrected keypoints showed significantly greater agreement with manual annotations than uncorrected data (Figure 1I).

Together, these components converged into two complementary pose estimation pipelines: an offline precision pipeline combining Mask R-CNN, DANNCE and SmoothNet for detailed kinematic analysis; and a real-time pipeline (YOLOv8 and DANNCE) operating at 35 fps while maintaining robustness to occlusion (Figure S1H, Video S3). The system’s accuracy scaled with camera count and remained high during high-occlusion social behaviors (Figures S1I-S1K). Fur pattern-specific training extended this capability to four simultaneously interacting animals (Figures S1L-S1M and Video S4), establishing a versatile and generalizable platform for studying naturalistic social behaviors.

### Feature-guided self-supervised clustering reveals juvenile social ethogram

With robust pose estimation achieved, we next asked whether unsupervised analysis of kinematic signatures could identify both known and novel social play motifs in juveniles. From 14 body keypoints, we derived 32 behavioral features spanning four categories: individual kinematics including speed, height, and flip angles; inter-animal spatial relations such as social distances and approach angles; sniffing engagement, defined as snout-to-body region overlaps; and body-occlusion dynamics during close contact (Figure 2A). Temporal autocorrelation analysis (Wiltschko et al., 2015) revealed that natural social interactions were organized at a characteristic timescale of ∼800 ms (_τ_ = 810 ± 177 ms, Figure 2B). We therefore segmented continuous social interactions into 800-ms action clips, yielding a dataset of 66,666 behavioral clips (800-ms each) from 40 juvenile pairs (PND 50-60; 20 male and 20 female pairs), focusing on contact-rich episodes (inter-animal distance <1 body length).

**Figure 2.**
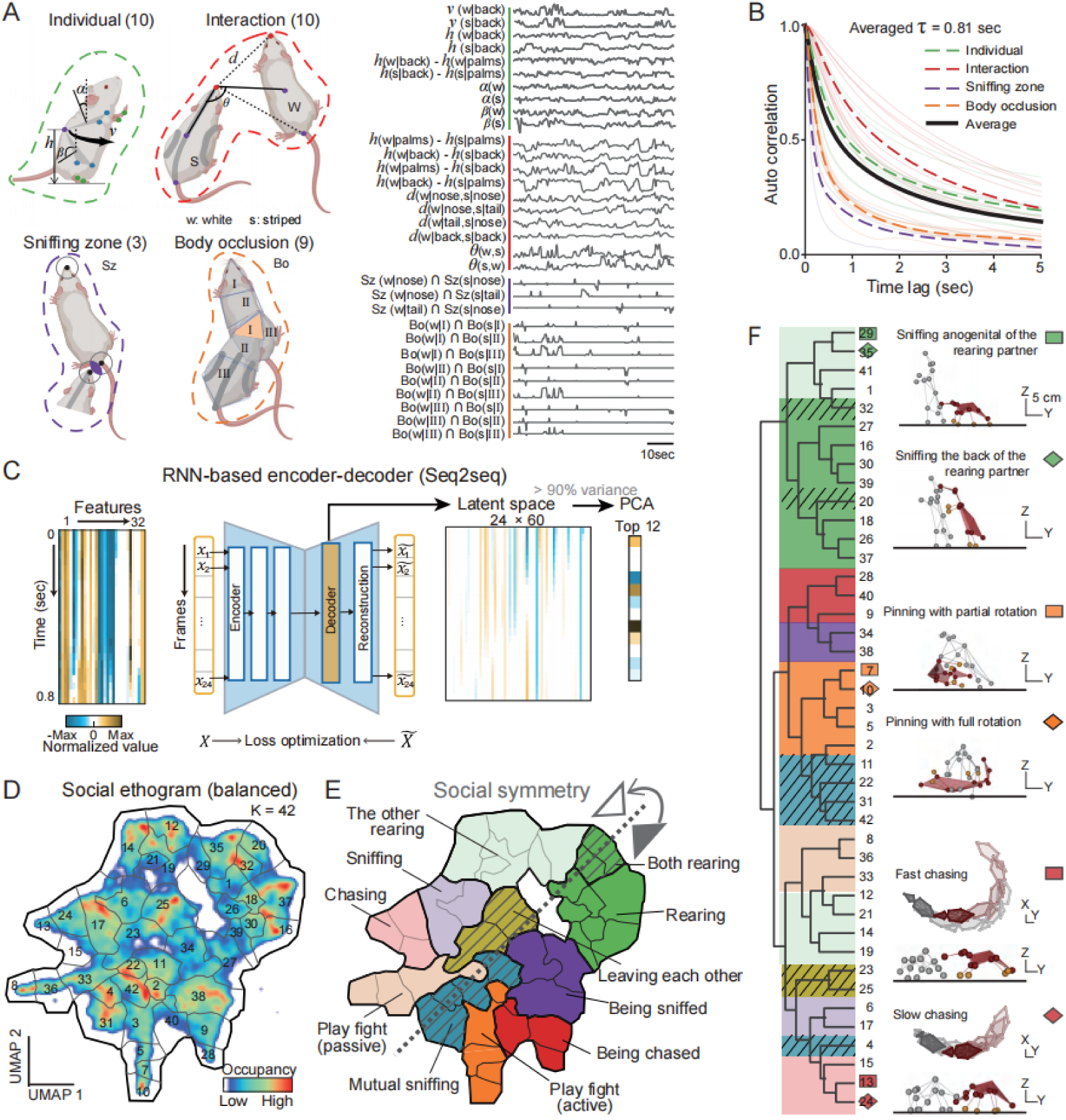
Self-supervised learning of a juvenile social interaction ethogram. (A) Social posture feature extraction. Quantified parameters included: Individual features: trunk velocity (*v*), back height (*h*), and upper/lower body plane angles (α*/*β); Interaction features: reciprocal nose-back-tail distances (*d*) and angles (θ), inter-animal height differences; Sniffing zones (Sz): overlap areas defined by nose-nose, nose-tail, and tail-nose regions (radii = average nose-ear/tail-claw distances; Body occlusions (Bo): XY-plane overlap areas among the three defined body regions (I: head; II: upper trunk; III: lower trunk), calculated for all pairwise combinations. (B) Temporal autocorrelations. Auto-correlograms of posture-related features declined and were fit with an exponentially decaying function to estimate the time-constant (_τ_). The averaged _τ_ was 0.81 s. (C) Seq2seq architecture. A three-layer, RNN-based encoder-decoder model processed 0.8-s clips (24 frames × 32 features) for self-prediction. The latent feature sequence (24×60) was reduced to 12 principal components (PCs, >90% variance explained). (D) Behavioral clustering. K-means (K = 42) clustering of 66,666 clips (subsampled from 86,560 clips across 40 rat pairs: 20 male/20 female) was visualized using UMAP with cluster boundaries overlaid. (E) UMAP projection reveals symmetric action patterns (dashed line), with color gradients indicating reciprocal behaviors (dark/light) and mutual interactions (slash). (F) Behavioral hierarchy. (Left) Dendrogram of cluster relationships based on pairwise Euclidean distances between cluster centroids in the latent space (12 PCs, STAR Methods). Branch colors match panel (E). (Right) Representative 3D skeletons from rearing, pinning, and chasing clusters, demonstrating that pose similarity correlates with dendrogram proximity. See also Figure S2.

To identify discrete interaction categories from these behavioral features, we trained a Seq2seq recurrent neural network (RNN)—an encoder-decoder architecture that captures spatiotemporal dynamics (Su and Shlizerman, 2020). PCA on latent features (top 12 PCs, 90% variance) revealed a high-dimensional ethogram partitioned into 42 behavior clusters by identifying local probability maxima (Figures 2C-2D and S2A). Strikingly, clusters self-organized into ethological categories: sniffing, chasing, pouncing, and leaving, among others, in an unsupervised manner (Figures 2E, S2B and Video S5). Crucially, reciprocal social roles were geometrically represented—symmetric positions on the UMAP manifold distinguished active *vs.* passive roles (e.g., sniffing *vs.* being sniffed; Figure 2E).

To quantify similarities and differences among clusters, we calculated cosine distances between latent features and used hierarchical clustering to group related behaviors. Such hierarchical clustering exposed micro-behavioral variants within broad categories (Figures 2F and S2C). For example, sniffing a rearing partner split into *anogenital vs. trunk* subtypes based on contact regions, pinning diverged by the pinned rat’s rotation angle (*partial vs. full*), and chasing split into *fast vs. slow* based on acceleration profiles. These granular distinctions, validated against expert ethologists’ annotations (Pellis et al., 2022), demonstrate our framework’s power to resolve subtle interaction mechanics—differences often lost in manual annotations.

### Cluster optimization through perspective-invariant labeling

While unsupervised clustering successfully identified ethologically relevant behaviors, substantial ambiguity remained at category boundaries—in which individual video clips often diverged from cluster prototypes, a well-documented limitation of unsupervised approaches (Hsu and Yttri, 2021; Tillmann et al., 2024). Given the intrinsic reciprocity of social interactions (Figure 2E), we posited that perspective-invariant labels (consistent across both participants’ viewpoints) would more cleanly demarcate high-confidence behavioral motifs.

To evaluate this, we established a cross-perspective consistency matrix by reprocessing each 800-ms clip from both rats’ viewpoints (Figures 3A and 3B). Clusters with > 50% inter-perspective overlap were considered reliable. Among these reliable clusters (Table S1), eight that showed identical labels for both rats were defined as mutual behaviors (e.g., cluster #4, head-to-head approach). The remaining clusters exhibited reciprocal labels, with active and passive roles forming complementary reciprocal role pairs (e.g., cluster #24 and #9, slow chasing *vs.* being slowly chased). Notably, some reliable clusters revealed over-or under-segmentation. For example, Clusters #5 and #7 mapped to the same reliable cluster (#36), indicating partial pinning behavior. In contrast, the ambiguous Cluster #14 split into two distinct clusters (#16 and #30), corresponding to distinct rearing types. These observations directly informed our subsequent cluster refinements (Figure 3C).

**Figure 3.**
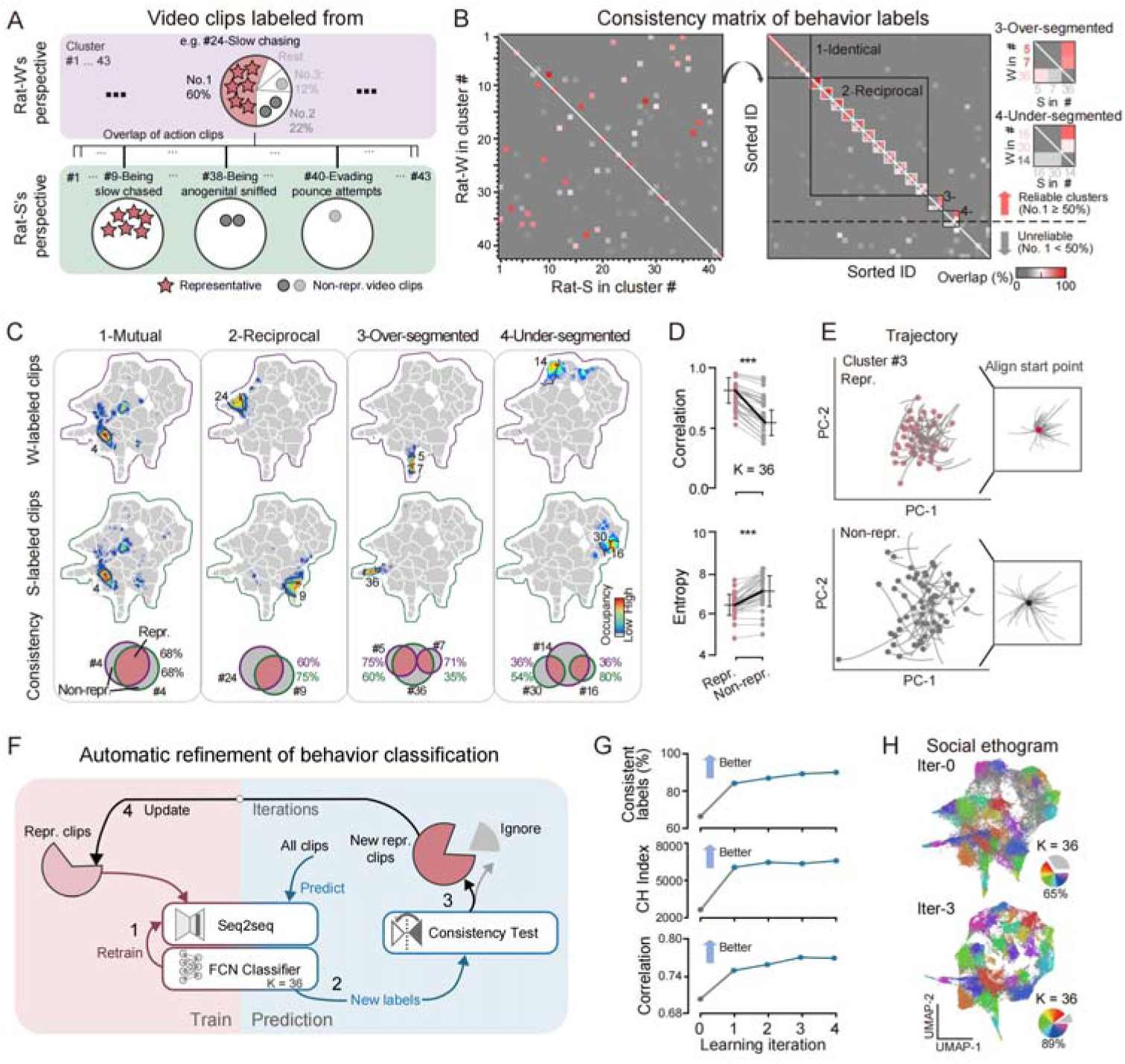
Active learning-based refinement of behavioral clusters. (A) Cross-perspective validation. Video clips from white (w) and striped (s) rat perspectives were compared within each cluster (e.g., cluster #24, slow chasing). Clusters with >50% inter-perspective overlap were classified as reliable, and their clips were considered representative. (B) Cross-perspective consistency matrix. Pairwise overlap percentages between w-and s-labeled clips across all clusters revealed 36 of 42 clusters as reliable (>50% overlap). Four relationship types emerged: (1) Mutual (8 clusters), showing high w-s overlap; (2) Reciprocal (12 clusters), matching a reciprocal cluster; (3) Over-segmented (clusters #5, #7), both mapping to a single reliable cluster (#36); (4) Under-segmented (cluster #14), mapping to multiple distinct clusters (#16, #30). (C) UMAP projections of w-labeled (top) and s-labeled (bottom) clips for representative cluster pairs illustrated in panel (B). Venn diagrams quantify clip overlap (pink) between perspectives (w: purple; s: green). (D) Comparison of intra-cluster correlation (top) and entropy (bottom) for representative (pink) vs. non-representative (grey) clips (\*\*\**p* < 0.001, two-sided paired t test). (E) Decoder trajectories in PC space. Dots indicate starting positions; insets show aligned trajectories. (F) Active learning pipeline. A Seq2seq-FCN framework iteratively improves labeling through: (1) supervised retraining of the Seq2seq-FCN on high-confidence clips; (2) prediction of new labels for all clips; (3) identification of reliable samples via cross-perspective consistency tests; (4) updating the high-confidence set with reliable samples for the next iteration. (G) Performance metrics across iterations. Representative clip proportion (top), Calinski-Harabasz Index (CHI; middle), and average intra-cluster correlation (bottom) plateaued after four iterations. (H) UMAP visualization of clip labels before (iter-0, top) and after refinement (iter-3, bottom). Representative (colored) and non-representative (grey) clips are shown, with pie charts indicating the proportion of representative clips. See also Figure S3.

Within reliable clusters, behavioral clips consistently labeled across perspectives were designated representative (Figure 3A). Representative clips within reliable clusters (64.8 ± 3.7% of total) exhibited higher pairwise latent feature correlations and lower entropy than non-representative clips (Figures 3D and S3A-S3B). In addition, temporal trajectories of representative clips were less dispersed in the latent space, further underscoring their reliability in capturing spatiotemporal dynamics (Figures 3E and S3C).

To automate cluster refinement, we first trained a fully connected classifier network (FCN) on high-consistency representative clips and then iteratively updated the Seq2seq latent space with FCN-predicted behavioral labels (Figure 3F, STAR Methods). Within four iterations, 89.2% of proximal-interaction clips (inter-animal distance < 1 body length) achieved perspective-invariant classification (*vs.* 64.8% initially; Figures 3G and 3H). Cluster validity indices improved significantly, as indicated by a 247% increase in the Calinski-Harabasz Index (CHI) and a 10.9% increase in inter-cluster correlation (Figure 3G). This final Seq2seq-FCN ethogram comprised 36 reliable behavioral clusters that generalized to distal interactions (Figures 3H, S3D-S3E). This optimized framework enables deep dissection of social behavioral dynamics across sexes and contexts.

### Dopaminergic encoding of sexually dimorphic social strategies

We began by characterizing naturalistic social interactions in same-sex juvenile rats (PND 50-60), identifying fundamental sex differences in behavioral strategies. Male pairs exhibited frequent contact-rich interactions dominated by investigatory sniffing followed by pouncing/pinning—a pattern characteristic of proactive social play. In contrast, female pairs adopted a social avoidance strategy, characterized by increased chasing and leaving bouts, along with heightened environmental exploration through rearing (Figures 4A-4C and S2D-S2J). These sex-specific strategies were further evident in action-sequence architecture, suggesting distinct reward values across social behavior categories (Figure 4C).

**Figure 4.**
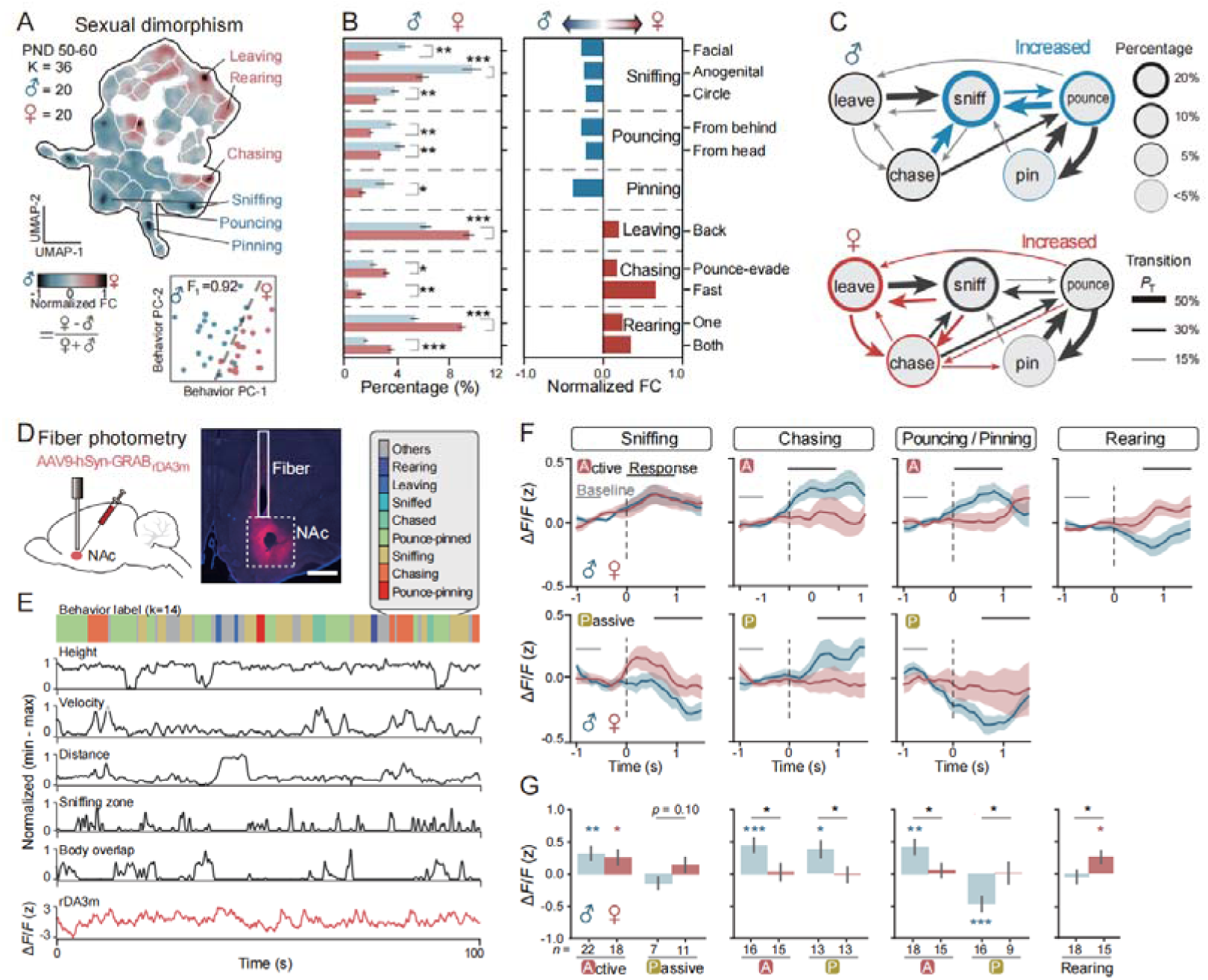
Sexually dimorphic behavioral and dopaminergic signatures in juvenile social interactions. (A) Sex differences in behavioral space. Top: Smoothed fold differences in UMAP cluster occupancy between male (blue; n = 20) and female (red; n = 20) rats. Bottom: Behavioral organization projected in low-dimensional space. The SVM classification boundary (dashed line; F1 score estimated by leave-one-out cross-validation; STAR Methods) separates sex-specific behavioral patterns. (B) Behavioral dimorphism. Left: Cluster proportions by sex. Right: Fold differences for semantically grouped behaviors (sniffing, pouncing, pinning, leaving, chasing, rearing; FDR-adjusted two-sided unpaired t test). (C) Behavioral state transitions. Averaged transition probabilities (*P_T_*) and cluster percentages for male (top) *vs.* female (bottom) pairs. Significant sex differences (*p* < 0.05, FDR-adjusted two-sided unpaired t test; STAR Methods) are color-coded. (D) Viral expression and fiber placement. Left: Sagittal AAV injection schematic. Right: Coronal section confirming GRAB-DA3m expression (red) and NAc core fiber placement (scale bar: 1 mm). (E) Representative DA recording. Time-aligned behavioral labels (top), kinematic variables (black), and z-scored ΔF/F (red) from a male pair. (F) DA dynamics during behavioral transitions. Z-scored ΔF/F aligned to behavioral onset (t = 0; STAR Methods). Traces shown mean ± SEM (male: blue; female: red; ≥3 epochs/video). Gray line: baseline (−1 to 0.5 s). Black line: response window (active: 0 to 1 s; passive: 0.5 to 1.5 s). (G)Peak z-scored ΔF/F (response window *vs.* baseline). Colored asterisks: within-group significance (two-sided paired t test); black asterisks: sex differences (two-sided unpaired t test). All error bars: mean ± SEM. \**p* < 0.05, \*\**p* < 0.01, \*\*\**p* < 0.001. See also Figure S4.

To investigate the neural basis of these differences, we expressed the dopamine sensor GRAB_DA3m_ in the NAc core and monitored dopamine fluctuations via fiber photometry during juvenile social interactions (23 male and 18 female pairs). Combining these recordings with our Seq2seq-FCN behavioral classification framework (36 clusters, 87.2% cross-perspective consistency; Figure S4A), we uncovered a tight coupling between NAc dopamine fluctuations and behavioral features across defined behavior types. Notably, DA responses tracked running velocity during chasing episodes, while during pouncing/pinning, DA responses correlated with the degree of body occlusion (Figures 4E, S4B-S4E).

Feature-aligned analysis (Figures S4B and S4F, STAR Methods) revealed sexual dimorphism in dopaminergic encoding (Figure 4F). During play behaviors, males exhibited elevated DA responses when actively pouncing/pinning (ΔF/F = 0.42 ± 0.11) but significant suppression when being pounced upon (ΔF/F = −0.47 ± 0.11), whereas this valence pattern was not observed in females (Figure 4F). This bidirectional signaling may reinforce active play initiation while discouraging passive states, explaining male-biased pouncing frequency. Conversely, females exhibited heightened DA responses selectively during non-social behaviors (e.g., rearing, ΔF/F = 0.26 ± 0.09), consistent with their predominant social avoidance strategy.

Surprisingly, some behavioral preferences were decoupled from the magnitude of dopamine responses. While females engaged in more chasing overall, males showed larger dopamine responses during active chasing (males: ΔF/F = 0.45 ± 0.11 *vs.* females: ΔF/F = 0.03 ± 0.13). Intriguingly, males exhibited stronger DA responses during passive chasing (being chased) (males: ΔF/F = 0.38 ± 0.13 *vs.* females: ΔF/F = −0.01 ± 0.11). A similar pattern emerged during sniffing interactions: although females initiated fewer sniffing episodes, they exhibited a trend-level increase in dopaminergic responses when being sniffed (males: ΔF/F = −0.14 ± 0.10; females: ΔF/F = 0.14 ± 0.12, p = 0.10). These results suggest that elevated DA responses during passive social engagement may potentially contribute to reduced initiation of active social interactions.

Collectively, our findings establish NAc dopamine as a sexually dimorphic encoder of social valence during juvenile development, with response patterns that both reflect and potentially shape sex-specific behavioral adaptations.

### Developmental trajectories of peer interactions in Shank3-deficient rats

Given our finding that NAc-DA dynamically encodes sex-specific social strategies, we asked whether disruption of these dynamics underlies the social motivation deficits that are central to ASD. To address this, we applied our integrated behavioral-photometry platform to *Shank3*-deficient (*Shank3*^+/−^) rats (exons 11-21 deletion; Meng et al., 2022) to determine how social interaction patterns diverge across developmental stages (Figure 5) and whether behavioral deficits during the juvenile period correlate with dysregulated DA dynamics (Figure 6).

**Figure 5.**
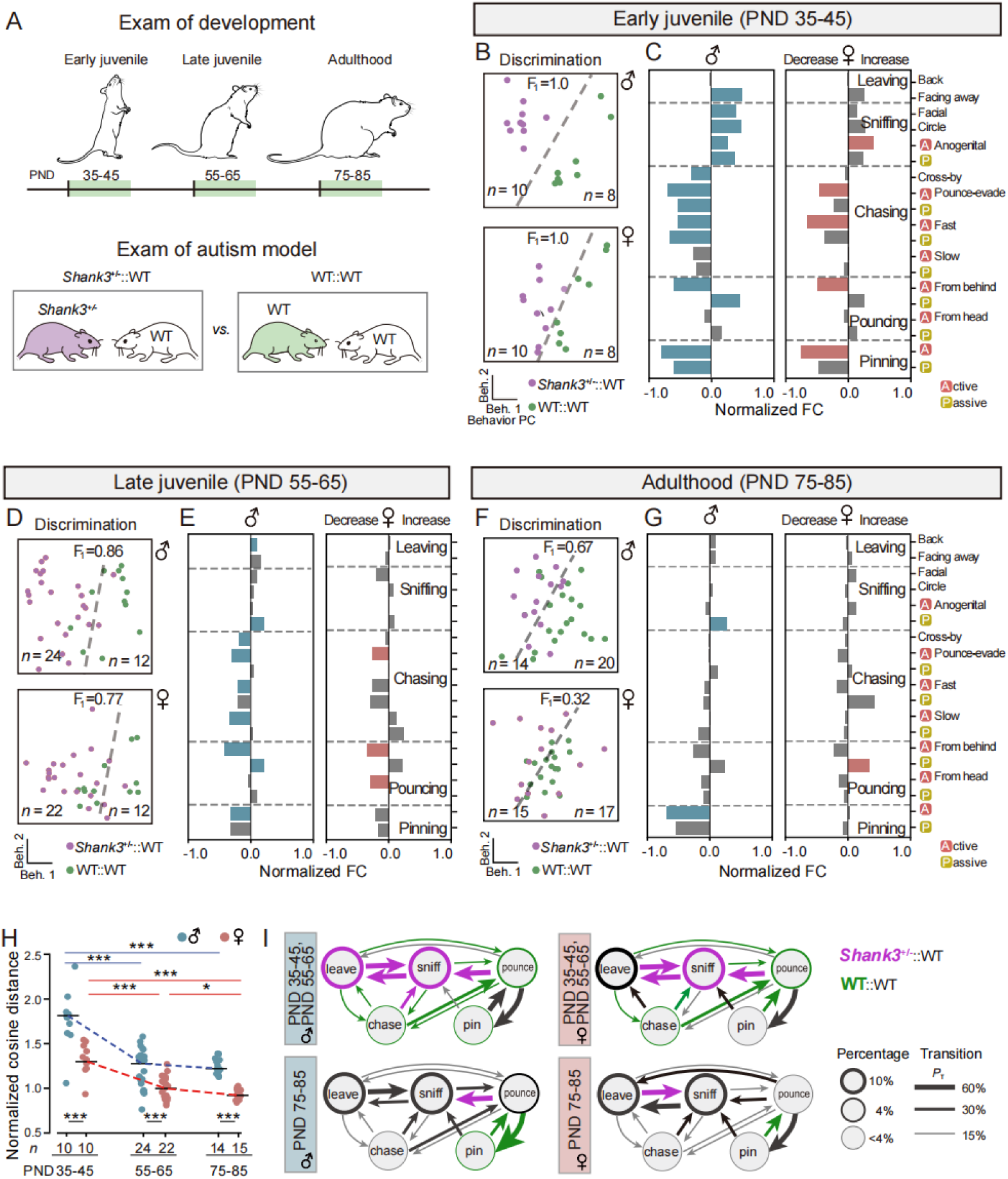
Age-and sex-dependent social behavior deficits in Shank3-deficient rats. (A) Experimental timeline. Freely moving social interaction tests were conducted across three developmental stages: early juvenile, late juvenile, and adulthood. (B, D and F) Behavioral profile clustering. PCA reveals distinct separation between *Shank3*^+/−^::WT (purple) and WT::WT (green) groups in males (top) and females (bottom). The SVM classifier boundary (dashed line; F1 score via leave-one-out cross-validation; STAR Methods) separates genotype-specific behavioral profiles. (C, E and G) Developmental behavioral alterations. Significantly altered social behavior categories (FDR-adjusted two-sided unpaired t test, *p* < 0.05) are color-coded (left: male; right: female). (H) Behavioral profile divergence. Cosine distance between *Shank3*^+/−^::WT and WT::WT groups, normalized to intra-group distances (males: blue; females: red; two-sided paired t test, \**p* < 0.05, \*\**p* < 0.01, \*\*\**p* < 0.001). (I) Behavioral state transitions. Comparison of juvenile (top) *vs.* adult (bottom) *Shank3*^+/−^rats (left: male; right: female). Average transition probabilities (*P_T_*) and cluster percentages for different genotype pairs. Significant genotype differences (FDR-adjusted two-sided unpaired t test, \**p* < 0.05) are color-coded. See also Figure S5.

**Figure 6.**
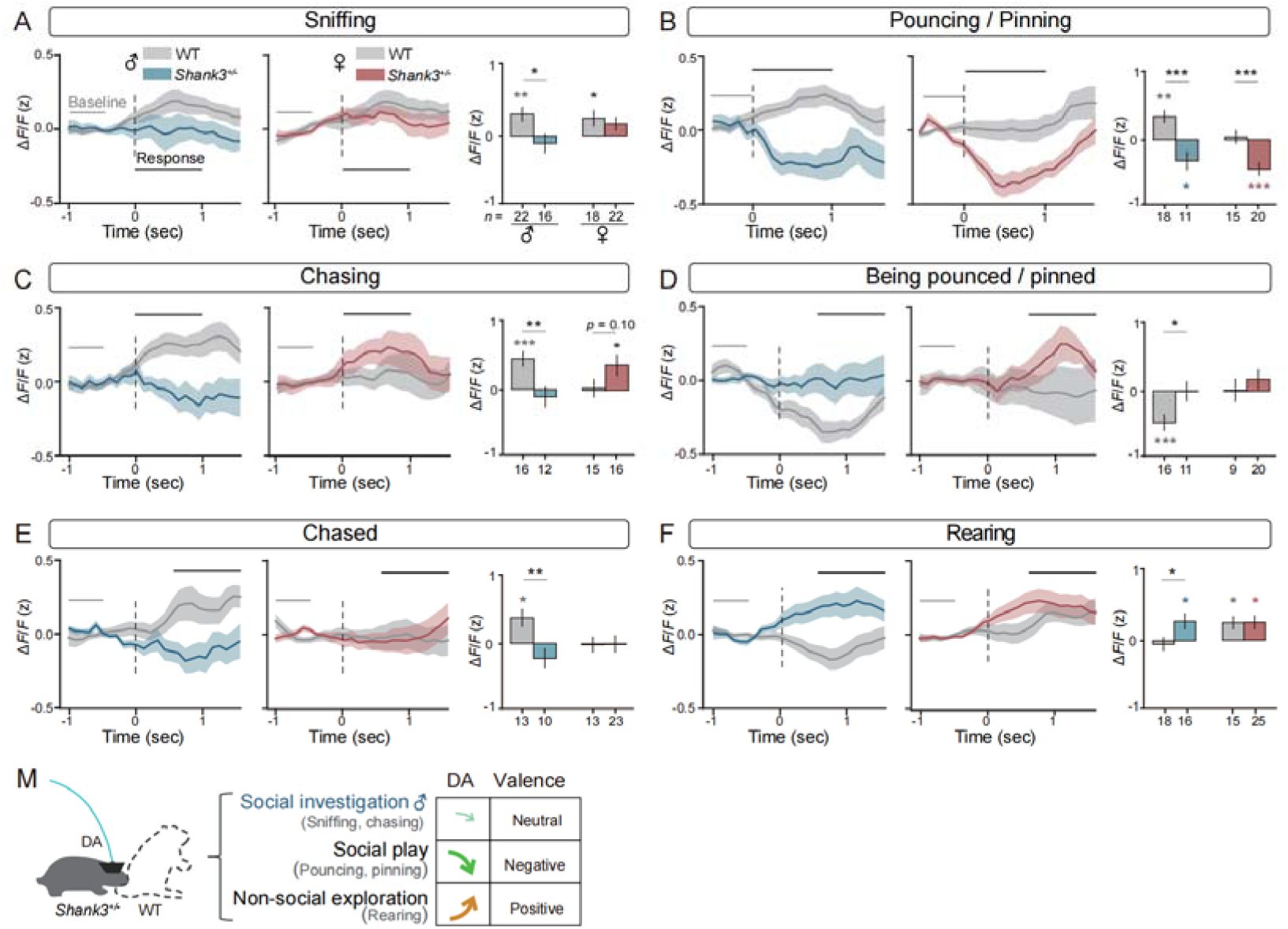
Dopaminergic responses across different behavioral categories in WT and Shank3+/−juvenile rats. (A-F) Time-resolved dopaminergic dynamics across different behavioral categories. Left and middle (traces): Z-scored ΔF/F aligned to behavioral onset (t = 0; STAR Methods), shown as mean ± SEM (male: blue; female: red; ≥ 3 epochs/video). Gray line: baseline (−1 to 0.5 s, −0.5 to 0 s in B). Black line: response window (active: 0 to 1 s; passive: 0.5 to 1.5 s). Right (bar plot): Peak z-scored ΔF/F (response window *vs.* baseline). Colored asterisks: within-genotype significance (two-sided paired t test); black asterisks: genotype differences (unpaired t test). (G) Summary of dopaminergic dysregulation in *Shank3*^+/−^juvenile rats. Behavioral categories with male-biased deficits are highlighted in blue. Error bars: mean ± SEM. \**p* < 0.05, \*\**p* < 0.01, \*\*\**p* < 0.001. See also Figure S6.

Longitudinal tracking of *Shank3*-deficient rats interacting with wildtype (WT) peers revealed age-dependent attenuation of social interaction deficits (Figures 5B-5G, S5A-S5B). At the early juvenile stage (PND 35-45), *Shank3*-deficient rats displayed hyperactive investigatory sniffing—sniffing bouts surged to 232% in males and 174% in females relative to WT levels—yet showed a dramatic failure to initiate play (pin-pouncing frequency decreased by 62% in males and 57% in females; Figure 5C). By the late juvenile stage (PND 55-65), play deficits attenuated yet persisted (pin-pouncing decreased by 33% in males and 44% in females; Figure 5E). However, in early adulthood (PND 75-85), *Shank3*-deficient males retained selective deficits in pinning (reduced by 83%), whereas *Shank3-*deficient females displayed normalized behaviors (Figure 5G).

Behavioral space analysis quantified these divergences: the distance that separated *Shank3*^+/−^and WT groups peaked during the juvenile stage and partially resolving by adulthood, with males showing greater deficits (Figures 5H and S5C-S5D). Strikingly, although WT partners adapted by increasing active play toward *Shank3^+/−^*rats (Figure S5E), *Shank3^+/−^*juveniles failed to sustain engagement—their play initiations more often devolved into non-progressive sniffing (Figure 5I). These behavioral trajectories suggest that *Shank3*-deficient juvenile rats retain a global interest in social investigation but lack the motivational drive required to execute coordinated play—a hypothesis we next tested using dopaminergic interrogation.

### Social reward deficits in Shank3-deficient juveniles

To examine the neural mechanisms underlying social motivation deficits in ASD, we used fiber photometry to record NAc DA dynamics in freely behaving *Shank3^+/−^*juveniles during social interactions with WT peers. Our recordings revealed profound dysregulation of reward-circuit function across discrete behavioral categories.

During social investigation, WT juveniles exhibited robust DA increases in approximately 40% of active sniffing events (male: ΔF/F = 0.32 ± 0.10; female: ΔF/F = 0.26 ± 0.11). In contrast, *Shank3^+/−^*juveniles showed markedly attenuated responses in males (ΔF/F = −0.11 ± 0.14) but not in females (ΔF/F = 0.18 ± 0.09, Figures 6A and S6A). No genotype differences were observed during passive sniffing (Figure S6G and S6H). This blunted signaling extended to chasing behaviors (both active and passive) specifically in males, with *Shank3^+/−^*juveniles failing to develop the characteristic positive DA responses seen in WT controls (Figures 6B-6C, S6B-S6C).

The most pronounced dysregulation emerged during social play (Figures 6D and S6D). WT males exhibited predominantly positive DA responses to pouncing events (ΔF/F = 0.35 ± 0.09), with 33% of events showing increased activity versus 28% showing decreased activity. In stark contrast, *Shank3^+/−^*juveniles displayed a complete valence reversal (ΔF/F = −0.34 ± 0.14), with only 21% of the events eliciting DA increases and 40% showing suppression. This pathological pattern was conserved in females (WT: ΔF/F = 0.04 ± 0.10, 38% increase *vs.* 39% decrease; *Shank3*^+/−^:ΔF/F = −0.45 ± 0.09, 22% increase *vs.* 46% decrease), suggesting that social play initiation is encoded with negative valence in *Shank3* mutants. Intriguingly, *Shank3*^+/−^males additionally lacked the characteristic DA suppression during passive play observed in WT males (WT: ΔF/F = −0.47 ± 0.11; *Shank3*^+/−^: ΔF/F = −0.01 ± 0.14; Figures 6E and S6E), revealing bidirectional impairment in both initiating and receiving play behaviors.

In contrast to social contexts, non-social exploration elicited significantly enhanced DA responses in *Shank3*^+/−^juveniles. During rearing, 53% of events in mutant males elicited positive DA responses (WT: ΔF/F = −0.05 ± 0.10; *Shank3*^+/−^: ΔF/F = 0.29 ± 0.11; Figures 6F and S6F). This contrasted sharply with WT males, who displayed balanced response profiles (35% increase *vs.* 39% decrease). A similar amplification pattern emerged in *Shank3*^+/−^females, with 49% of events showing increased DA activity compared to 27% decreases, while WT females maintained equilibrium (38% increase *vs.* 35% decrease, Figure S6F). However, the peak response amplitude did not differ significantly between female genotypes (Figure 6F).

Together, these results demonstrate comprehensive dysregulation of valence coding in *Shank3* deficiency, characterized by blunted DA signaling during social investigation, pathological negative valence assignment to active social play, and enhanced reward valuation of non-social exploration. The more pronounced DA dysregulation in males paralleled their more severe behavioral deficits (Figure 5), providing a potential neural mechanism underlying the male predominance in ASD social impairments.

### Closed-loop activation of VTA-NAc pathway in *Shank3*^+/−^juveniles

Our fiber photometry recordings revealed disrupted DA dynamics in the NAc core during social interactions in *Shank3*^+/−^juveniles, which correlated with specific play-behavior impairments. To establish causality, we developed a closed-loop optogenetic system that selectively activated tyrosine hydroxylase (TH)-positive VTA DA terminals in the NAc core during impaired social behaviors (Figures 7B-7D). This intervention was administered during daily 15-minute naturalistic interactions with WT partners over eight consecutive days (PND 46-54), followed by a 10-day post-stimulation observation period (Figure 7A).

**Figure 7.**
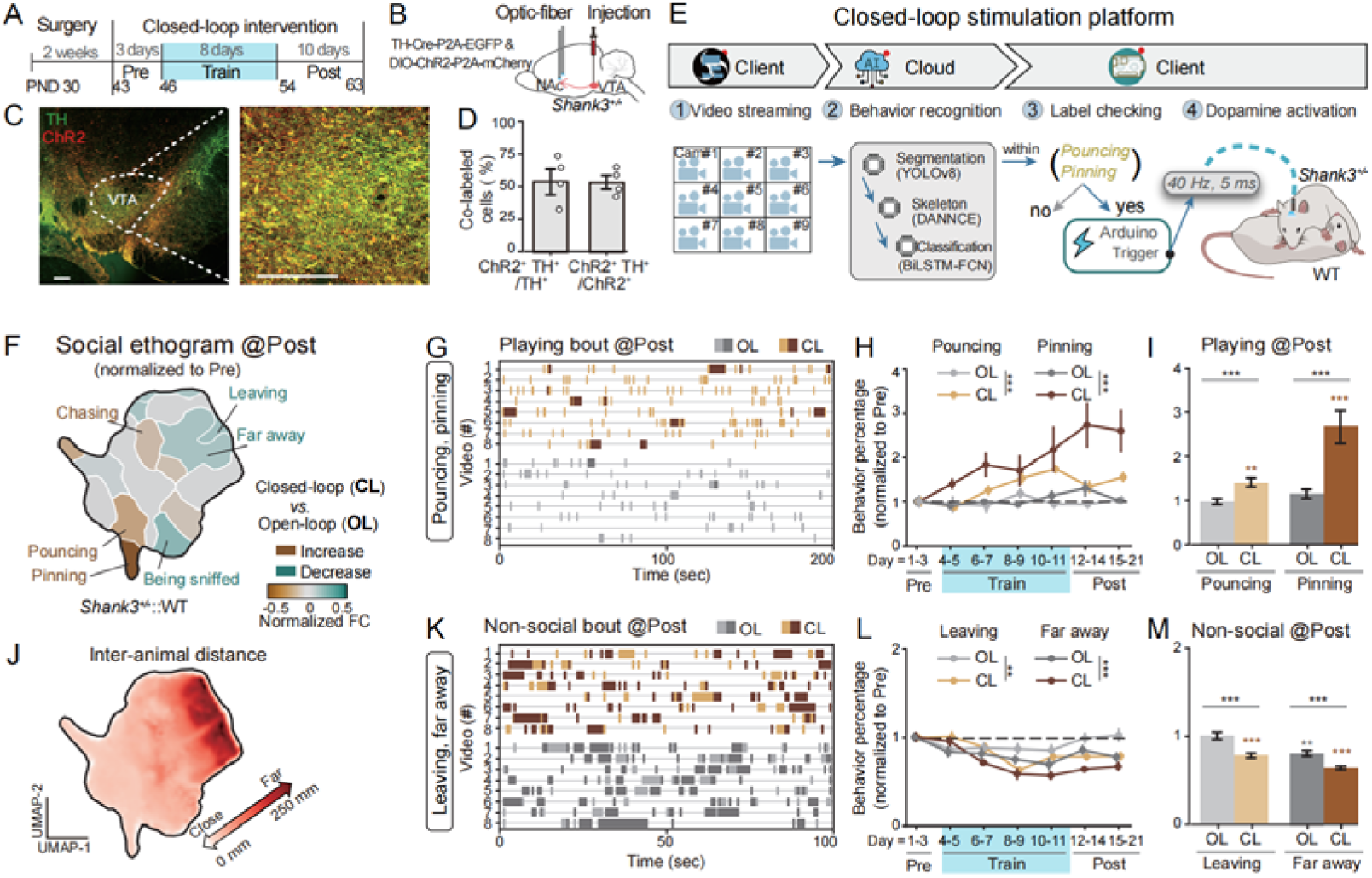
Closed-loop NAc-DA activation rescues social play deficits in Shank3+/−juvenile rats. (A) Experimental timeline. The 8-day closed-loop optogenetic intervention and 10-day post-stimulation assessment were performed in *Shank3*-deficient males (closed-loop: n = 9; open-loop: n = 5; PND 43-63). (B) Viral targeting strategy. TH-Cre-driven ChR2-mCherry expression in VTA dopamine neurons with optical fiber implantation in the NAc core. (C) Histological verification. ChR2-mCherry (red) colocalization with TH^+^ neurons (green) in the VTA. Scale bars: 200 _μ_m. (D) Quantification of ChR2^+^ and TH^+^ co-expression in VTA (n = 4 rats). (E) Closed-loop stimulation system. Multi-camera inputs were processed via YOLOv8-based segmentation, DANNCE-derived 3D pose estimation and BiLSTM-FCN classification (36 behavioral categories) in a cloud pipeline. Detected pouncing/pinning events triggered 40-Hz blue light pulses (5-ms pulse, 0.5-s duration) to NAc-projecting VTA terminals (<400 ms latency). (F) Post-training behavioral changes. UMAP cluster occupancy (normalized to pre-training baseline) showing increased (yellow)/decreased (cyan) behaviors in closed-loop (CL) *vs*. open-loop (OL) groups. (G-I) Play behavior analysis. (G) Representative raster plots: pouncing (light color) and pinning (dark color) in OL (gray) *vs*. CL (yellow) during the post-training period. (H) Progressive play duration changes (one-way ANOVA). (I) Total play behavior duration after training (CL: 32 videos; OL: 30 videos; two-sided unpaired t test). (J) Social proximity mapping. UMAP embedding color-coded by inter-animal distance (warmer colors indicate greater separation). (K-M) Non-social behavior comparison. Analysis of leaving (light color)/far-away (dark color) clusters (same conventions as in G-I). Error bars: mean ± SEM. \**p* < 0.05, \*\**p* < 0.01, \*\*\**p* < 0.001. See also Figure S7.

The behavioral detection system employed a custom cloud-based real-time analysis pipeline capable of processing multi-camera video streams, recognizing behavioral categories, and triggering optogenetic stimulation during target behaviors with an end-to-end latency of <400 ms (Figures 7E and Video S7). The pipeline sequentially processed each video frame, beginning with YOLOv8-based animal segmentation, followed by DANNCE-derived 3D pose estimation and feature extraction. These features were then classified into 36 behavioral categories by a bidirectional LSTM-Fully Convolutional Network (BiLSTM-FCN) (Figures S7A-S7C). Behavioral classification for each frame was determined by analyzing the preceding 400-ms behavioral sequence, achieving a mean F1 score of 0.8 across all behavioral categories (Figures S7D and S7E). Given the high autocorrelation of rat social behaviors within 800-ms windows (Figure 2A), this design ensured accurate capture of ongoing social interactions while maintaining the temporal precision required for closed-loop optogenetic intervention.

Using this system, we delivered closed-loop 40 Hz stimulation to NAc DA fibers specifically during deficient play behaviors (pouncing/pinning) (Figure S7F), which accounted for 12.1 ± 1.0% of total behavioral testing time and were characterized by aberrant negative DA dynamics in *Shank3*^+/−^juveniles (Figure 6D). Open-loop controls received temporally unpatterned stimulation matched in total duration (11.2 ± 0.3%; Figure S7G). Notably, closed-loop stimulation delivered 19.6 ± 1.8% of its pulses during active play initiation bouts, compared to only 0.7 ± 0.1% in open-loop controls (Figure S7I), while maintaining high behavioral recognition accuracy (>90% true positive rate for target behaviors, <15% false positive rate for non-target behaviors; Figure S7H).

Closed-loop activation induced persistent behavioral changes over consecutive training days and throughout the 10-day post-stimulation period (Figures 7F and S7J-S7K). The most pronounced stimulation-biased effects were observed in behavioral clusters corresponding to pouncing and pinning, which showed 1.4-fold and 2.7-fold increases in duration, respectively, compared to pre-stimulation baselines (Figures 7G-7I and S7L). Moreover, closed-loop-stimulated mutants exhibited a reduction in time spent in non-social behavioral clusters (e.g., leaving, far-away) that occupy spatially distinct regions in UMAP space and are characterized by larger inter-animal distances compared to other behavioral clusters (Figures 7J-7M).

Intriguingly, WT partners interacting with closed-loop-stimulated mutants also exhibited altered behavioral profiles, with increased pinning and chasing and decreased sniffing (Figure S7M), indicating global adaptive changes in social motivation that thereby promoted enhanced social engagement between *Shank3*^+/−^and WT dyads. Together, these results demonstrate that precisely timed DA release during play behaviors can effectively rescue social-circuit dysfunction in *Shank3*^+/−^juveniles, supporting the hypothesis that behavior-specific neuromodulation may ameliorate developmental social deficits.

## Discussion

By integrating ethologically relevant behaviors, dopaminergic dynamics, and circuit manipulation, our findings demonstrate that NAc dopamine acts as a sex-specific modulator of social motivation. Notably, we identify a paradoxical reversal of reward processing in *Shank3-*deficient models — where initiating social play is encoded with negative valence, whereas non-social exploration is disproportionately reinforcing — mirroring clinical observations of atypical reward processing in children with ASD (Cascio et al., 2012; Kim et al., 2014). These results not only elucidate the developmental neurobiology of social motivation but also provide a framework for targeted circuit interventions during social critical periods, offering new translational avenues for ASD therapeutics.

### Identification of juvenile-typical social ethogram

Our study introduces a high-throughput, low-latency behavioral tracking platform optimized for juvenile social play in rodents. While recent breakthroughs like social-DANNCE enabled multi-animal 3D pose estimation in adults (Klibaite et al., 2025), our nine-camera system provides superior resolution for juvenile-specific behaviors involving intense physical contact (e.g., pouncing, pinning) and rapid somersaults. Beyond hardware superiority, our cloud-accelerated pipeline achieves a maximum processing latency of 400 ms — a critical advance that enables real-time behavior identification synchronized with neural manipulation. This temporal precision permits closed-loop optogenetic interventions at specific behavioral transitions (e.g., pounce initiation), which were previously impossible with offline methods.

By combining ethologically relevant feature extraction with active learning using clips with cross-perspective-consistent labels, we addressed a persistent challenge in unsupervised behavioral clustering: the trade-off between interpretability and granularity (Jiao et al., 2022). Our iterative refinement approach achieved 87.2% cross-perspective consistency, producing an ethogram that aligns with expert annotations while retaining computational objectivity. This framework establishes a standardized behavioral atlas for juvenile social interactions, enabling precise quantification of individual differences across genotypes and developmental stages. Importantly, the modular design permits seamless adaptation to diverse social paradigms—from dyadic play to group interactions—offering a universal platform for studying social behaviors in different contexts.

### Dopaminergic encoding of social valence and its breakdown in *Shank3* deficiency

We identify NAc dopamine as a dynamic encoder of social valence that governs sex-and context-dependent behavioral strategies during development. In wild-type juveniles, dopaminergic signaling exhibited adaptive modulation during naturalistic interactions, with phasic increases during active social initiation (i.e., sniffing, chasing, pouncing/pinning), while passive engagement typically elicited neutral responses. However, critical exceptions revealed specialized reward processing mechanisms (Figures 4F and 4G): male juveniles showed strong DA release when being chased, while females exhibited increased DA responses when being sniffed, coinciding with their reduced initiation of these specific behaviors. Most strikingly, being pounced upon triggered pronounced DA suppression exclusively in males—a valence inversion that may reinforce their characteristically high frequency of pouncing/pinning initiation—whereas females lacked this response pattern. These dopaminergic signatures precisely mirror the observed behavioral strategies, with males engaging in frequent play but minimal chasing, and females showing the opposite preference.

Females further diverged through amplified DA responses during non-social exploration, particularly rearing, mirroring their preference for environmental sampling over sustained social contact. These sex-specific patterns may reflect evolutionary adaptations. Males utilize dopamine to reinforce playful fighting that establishes social hierarchies (Achterberg and Vanderschuren, 2023), while females may employ DA modulation to mitigate social over-engagement and reduce anxiety-related responses during prolonged interactions (Asher et al., 2017).

In *Shank3*-deficient rats, this adaptive flexibility collapses (Figures 6). Both sexes exhibit paradoxical suppression during play initiation, essentially abolishing their capacity for peer play. *Shank3*-deficient males additionally showed reduced DA increases during appetitive investigation, blunted DA suppression during passive play, and DA hyper-responsivity during rearing, mechanistically reducing social initiative while reinforcing non-social exploration. These deficits peak during the juvenile critical period, when young animals refine social skills through play, paralleling the trajectory of social impairment in ASD where early play deficits often precede lifelong social-cognitive challenges (Jordan, 2003). The striking parallels between these neural signatures and core ASD phenotypes highlight the importance of NAc dopamine dynamics in shaping social behavior strategies.

### Targeted intervention strategies

Our demonstration that closed-loop dopamine stimulation produces both acute and sustained rescue of social play deficits in *Shank3*-deficient juveniles reveals several fundamental insights into targeted neuromodulation. The persistence of behavioral improvements ten days after stimulation cessation (Figure S7D) suggests that transient, behaviorally patterned DA activation can drive lasting circuit reorganization during developmental critical periods. This durable effect aligns with established mechanisms of reinforcement learning, where phasic DA signals strengthen action-outcome associations (Markowitz et al., 2023) and extends these principles to social behavior remediation (Solié et al., 2022; Willmore et al., 2022). The critical period’s enhanced plasticity for social reward learning likely created optimal conditions for such DA-dependent circuit remodeling.

The therapeutic potential of this approach may be further enhanced by engaging complementary neuromodulatory systems. While our study focused on DA, other systems—particularly serotonin (5-HT) and oxytocin (OT) —are known to regulate social motivation during critical periods and exhibit disrupted signaling in ASD models (Dölen et al., 2013; Nardou et al., 2023; Walsh et al., 2021, 2018). These systems show complex synergies with DA in the NAc. For example, co-activation of DA and 5-HT enhances reinforcement more potently than either system alone (Cardozo Pinto et al., 2025), while OT potentiates DA-mediated social reward signaling (Rappeneau and Castillo Díaz, 2024). Our real-time behavioral platform is uniquely positioned to investigate these interactions, as its ability to precisely link neuromodulatory events to behavioral outcomes enables systematic evaluation of both immediate effects and long-term plasticity induced by combinatorial interventions. Future studies could leverage this system to identify optimal sequences or ratios of DA, 5-HT, and OT engagement to maximize durable social circuit reorganization.

## Resource availability

### Lead contact

Further information and requests for resources and reagents should be directed to and will be fulfilled by the lead contact, Ying Li.

### Data availability

All raw experimental data and analytical results generated in this study will be provided upon reasonable request.

### Code availability

All custom code used in this study is available on GitHub (https://lilab-cibr.github.io/Social_Seq/en/).

## Acknowledgements

The authors thank Q. Yang and X. Mi for technical assistance; CIBR LARC staff for animal care; CIBR Imaging Core, Instrumentation Core, and Vector Core for technical support; R. Zhang for providing the *Shank3*-mutant rat; Y.L Li for providing the dopamine sensor construct; W. Xiong and X. Yu for constructive comments on the manuscript; All members of the Li laboratory for helpful comments on this project. This work was supported by grants from: The Key Program of the National Natural Science Foundation of China (No. 32330045), STI2030-Major Projects (2021ZD0203903), CAMS Innovation Fund for Medical Sciences (2024-I2M-ZD-013), the Natural Science Foundation of Beijing, China (JQ24036), and Human Frontier Science Program, Career Development Award (CDA00005/2019). X.M.T. is supported by the fellowship of China Postdoctoral Science Foundation (2021M700454).

## Author Contributions

Conceptualization: Y.L., X.F.C. and X.M.T.; methodology: Y.L., X.F.C., X.M.T., Z.C.Z., M.W; formal analysis: X.F.C., X.M.T., Y.O., and Z.Y.D.; experiments: Z.C.Z., X.F.C., X.M.T., Y.Q.Z., Y.X.L., M.A. and M.W; writing -original draft: Y.L., X.F.C. and X.M.T.; visualization: X.F.C., X.M.T and Z.C.Z.; code release: X.F.C., X.M.T; supervision: Y.L.; funding acquisition: Y.L., and X.M.T.

## SUPPLEMENTAL FIGURES

**Figure S1.**
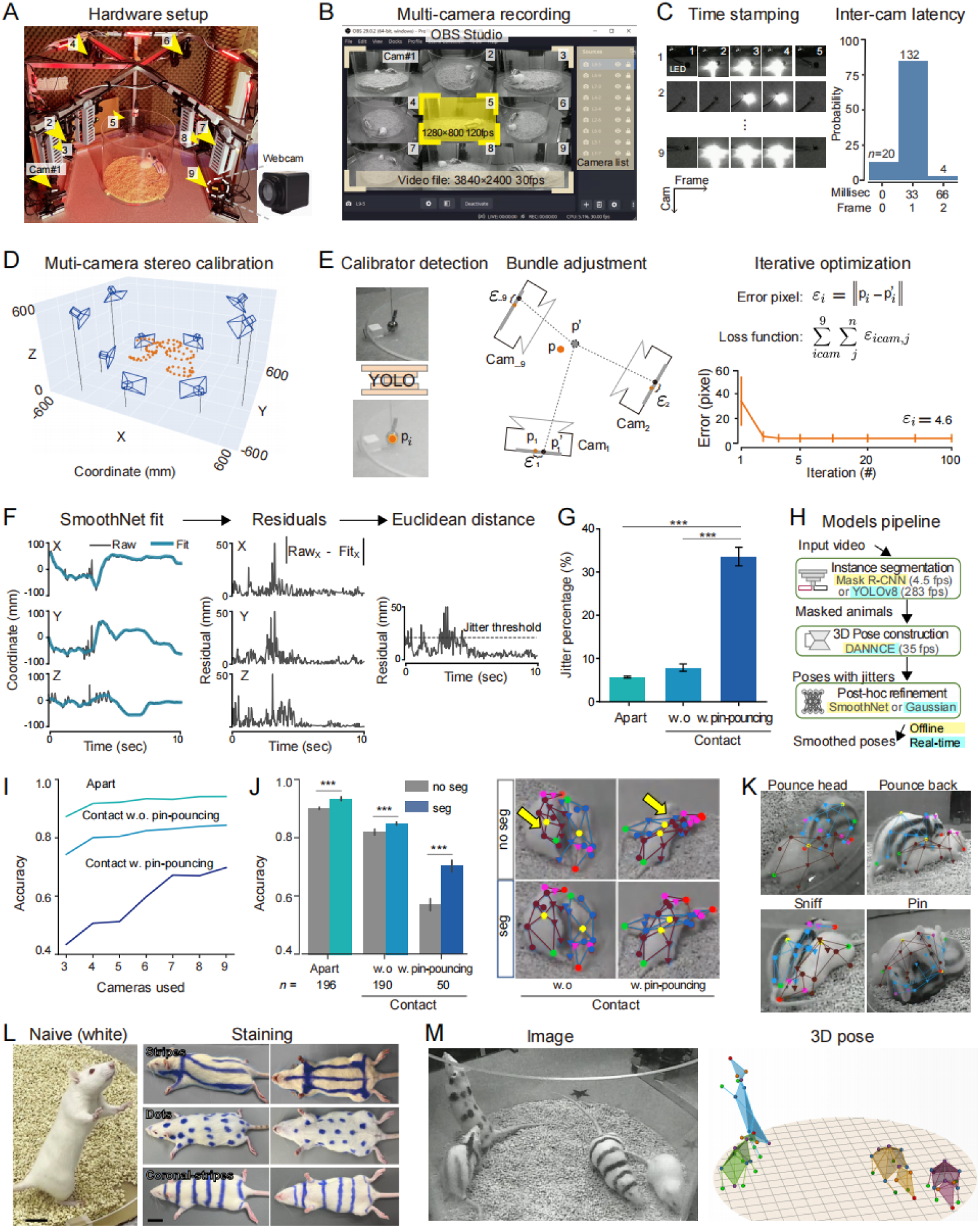
Experimental setup and 3D social pose reconstruction pipeline, Related to Figure 1. (A) Recording environment. Circular arena with nine synchronized webcams under red illumination. (B) Video acquisition. Videos captured by different cameras were arranged in a 3×3 grid mosaic at 3840×2400 resolution and were downsampled to 30 fps using OBS Studio. (C) Temporal synchronization. Left: LED pulse detection across cameras. Right: 95% of inter-camera latencies were ≤ 1 frame (33 ms). (D) Extrinsic parameter calibration for multiple cameras (blue) using a randomly moving spherical calibrator (orange) as the reference object (STAR Methods). (E) Spatial calibration workflow. Left: YOLOv8-based detection of the spherical calibrator. Middle: Initial camera alignment show that the 3D position of the calibrator (*p’*) deviates from the actual position (*p*), with corresponding pixel projection errors (_ε_*_i_*) on each camera plane (*cam_i_*). *Right:* Iterative optimization minimizes the projection error (_ε_*_i_*) across a set of calibrator points (n = 50). The line plot shows progressive error reduction across iterations (mean ± SD). (F) Motion jitter correction. Left: SmoothNet-derived kinematic fits (blue) superimposed on raw 3D keypoint trajectories (gray) along X/Y/Z axes. Middle, Residual displacements between raw and fitted positions for each spatial dimension. Right, 3D Euclidean residuals (>2-3 SD threshold; red: jitter events; STAR Methods). (G) Jitter percentage by social context. Apart (inter-animal distance ≥5 cm), Close contact (<5 cm), or Play (pinning/pouncing) (n = 8 videos; one-way ANOVA with Tukey post hoc test; mean ± SEM). (H) Pipeline comparison. Offline (yellow) *vs.* real-time (cyan) pipelines. (I) Landmark prediction accuracy. Improves with camera count but declines with occlusion severity (10-mm threshold). (J) Segmentation benefits. Left: Landmark prediction accuracy with (gray) or without (blue) prior segmentation (\*\*\**p* < 0.001, two-way ANOVA with Tukey post hoc, mean ± SEM). Right: Representative 3D predictions showing error reduction (yellow arrows). (K) Representative examples of accurate landmark predictions in high-contact social interactions (L) Fur patterning for individual identification during multi-animal tracking. Scale bars, 2 cm. (M) Multi-animal tracking. Simultaneous identity maintenance tracking for four rats.

**Figure S2.**
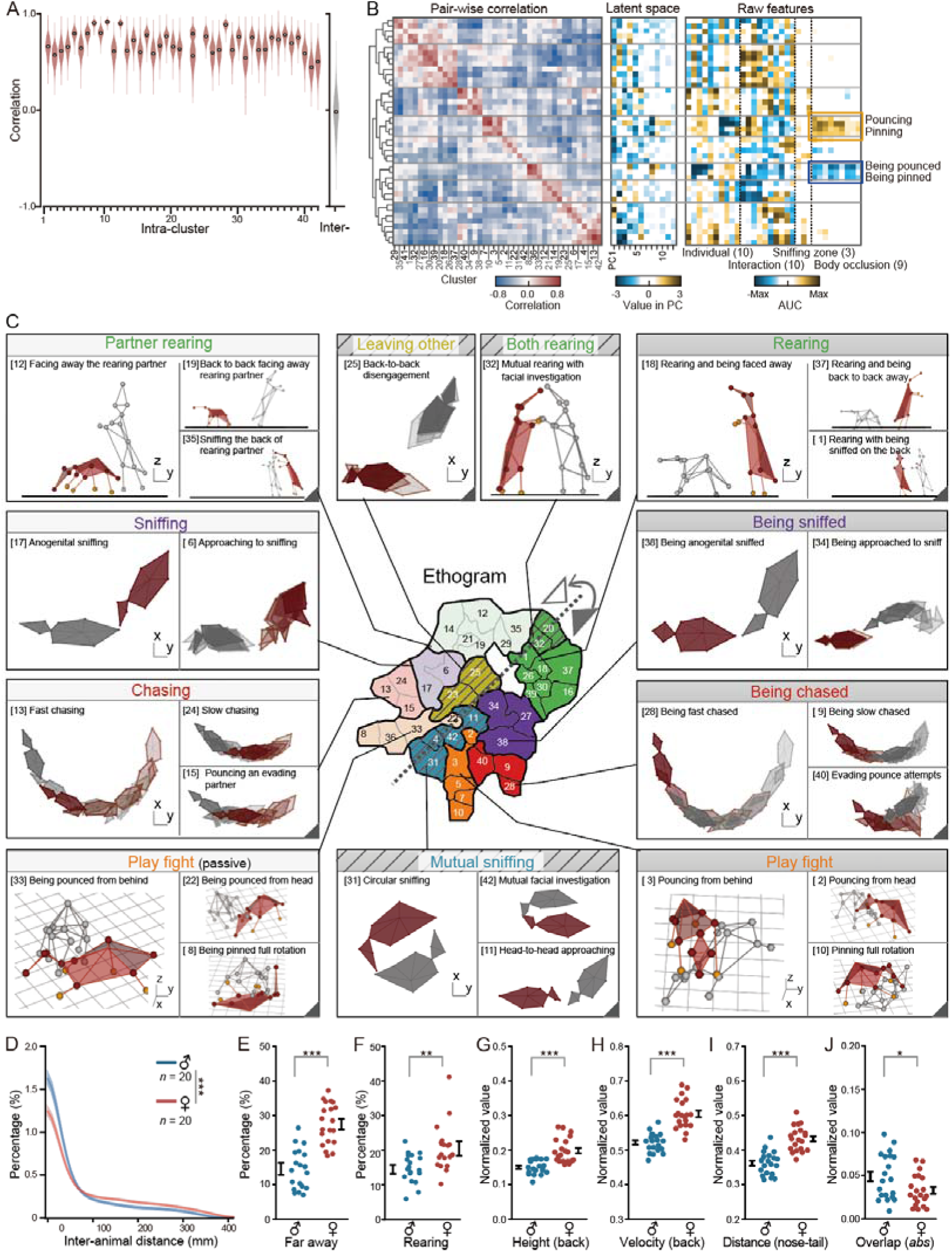
Social ethogram and sexually dimorphic behavioral features, Related to Figures 2 and 4. (A) Cluster validation. Violin plots of pairwise correlations of latent vectors (top 12 PCs). We compared intra-cluster (red) versus inter-cluster (gray) clip pairs (n = 200 random sampled pairs per distribution). (B) Behavioral modularity. Left: Correlation matrix of 42 behavior clusters based on latent vectors. The dendrogram is the same as in Figure 2F. Middle: Averaged latent vectors for each cluster (top 12 PCs). Right: Discriminant kinematic features (AUC values) for prototypical behaviors. Highlighted features include body overlap ratios in pouncing/pinning clusters. AUC: area under the curve. (C) Behavioral exemplars. Characteristic pose sequences representing each cluster in the social behavior atlas. (D) Social proximity. Inter-animal distance distributions for male (blue; n = 20 videos) vs. female (red; n = 20 videos) pairs. Calculated as the minimum XY-plane keypoint separation (mean ± SEM; Kolmogorov-Smirnov test). (E-J) Sex differences in social behaviors. (E) Time spent >1 body length apart. (F) Rearing duration. (G-J) Normalized behavioral features including height (G), velocity (H), nose-tail distance (I), and absolute body-overlap values (J). Data shown as mean ± SEM; two-sided paired t test; \**p* < 0.05, \*\**p* < 0.01, \*\*\**p* < 0.001.

**Figure S3.**
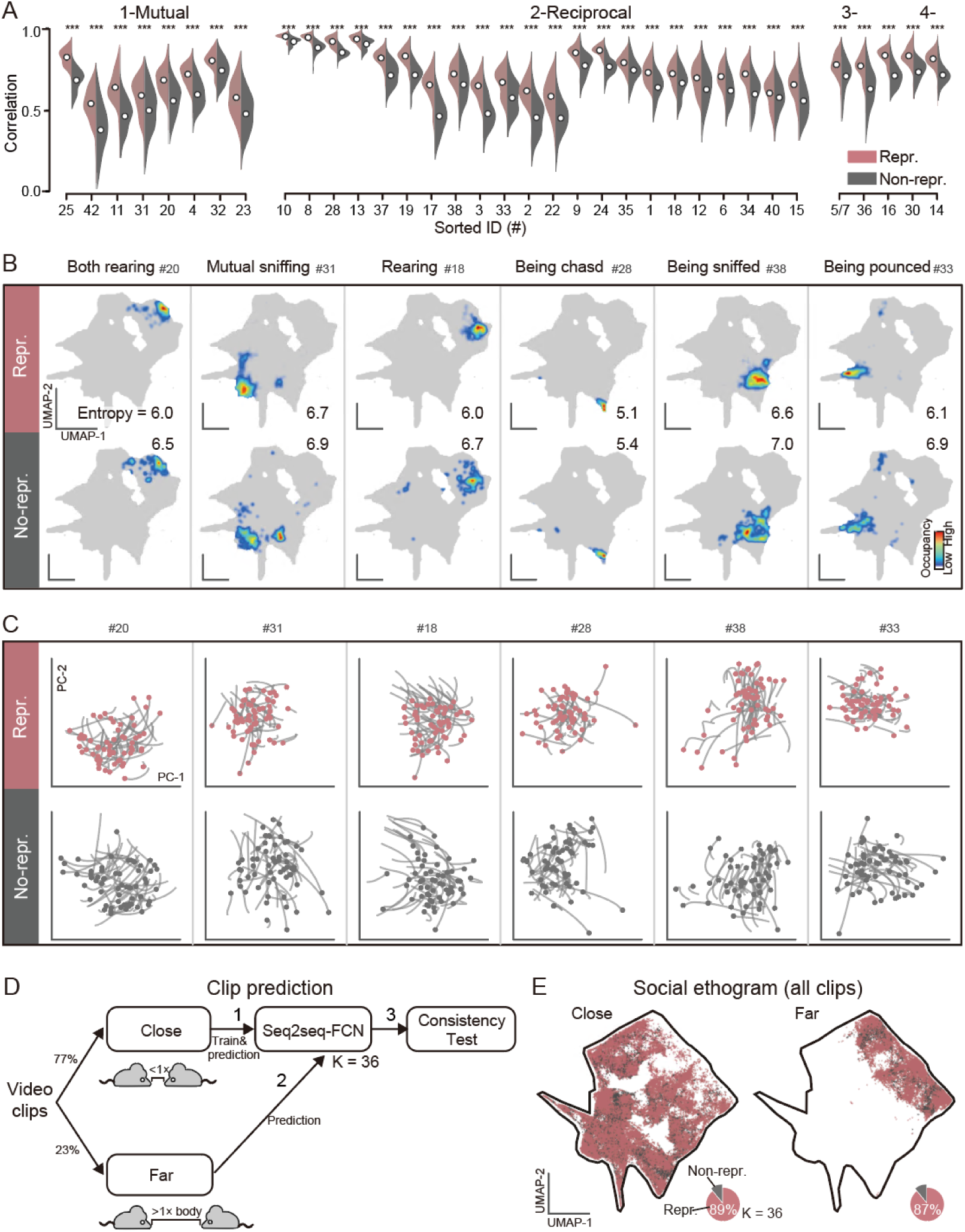
Cluster refinement and evaluation of representative behaviors, Related to Figure 3. (A) Cluster quality validation. Representative clips (red) show significantly higher intra-cluster correlation in latent space (top 12 PCs) compared to non-representative clips (gray; \*\*\**p* < 0.001, two-sided unpaired t test). Each distribution includes 200 randomly sampled clip pairs from reliable clusters. (B) UMAP visualization. Projection of representative *vs.* non-representative clips, with cluster entropy computed from spatial density estimates (STAR Methods). (C) Behavioral dynamics. Decoder trajectories in the top two PCs, with dots indicating starting points for individual clips. (D) Schematic of iterative refinement pipeline. (1) Training: the Seq2seq-FCN model is trained on high-confidence “Close” clips (inter-animal distance < 1 body length); (2) Application: the model generalizes to “Far” clips (distance > 1 body length; K = 36 clusters). (3) Selection: representative clips are identified through cross-perspective consistency testing. (E) Refinement outcomes. Behavioral-space distribution of training (“Close”, left) and application (“Far”, right) datasets. Representative (red) and non-representative (gray) clips are shown, with pie charts indicating the proportions of representative clips.

**Figure S4.**
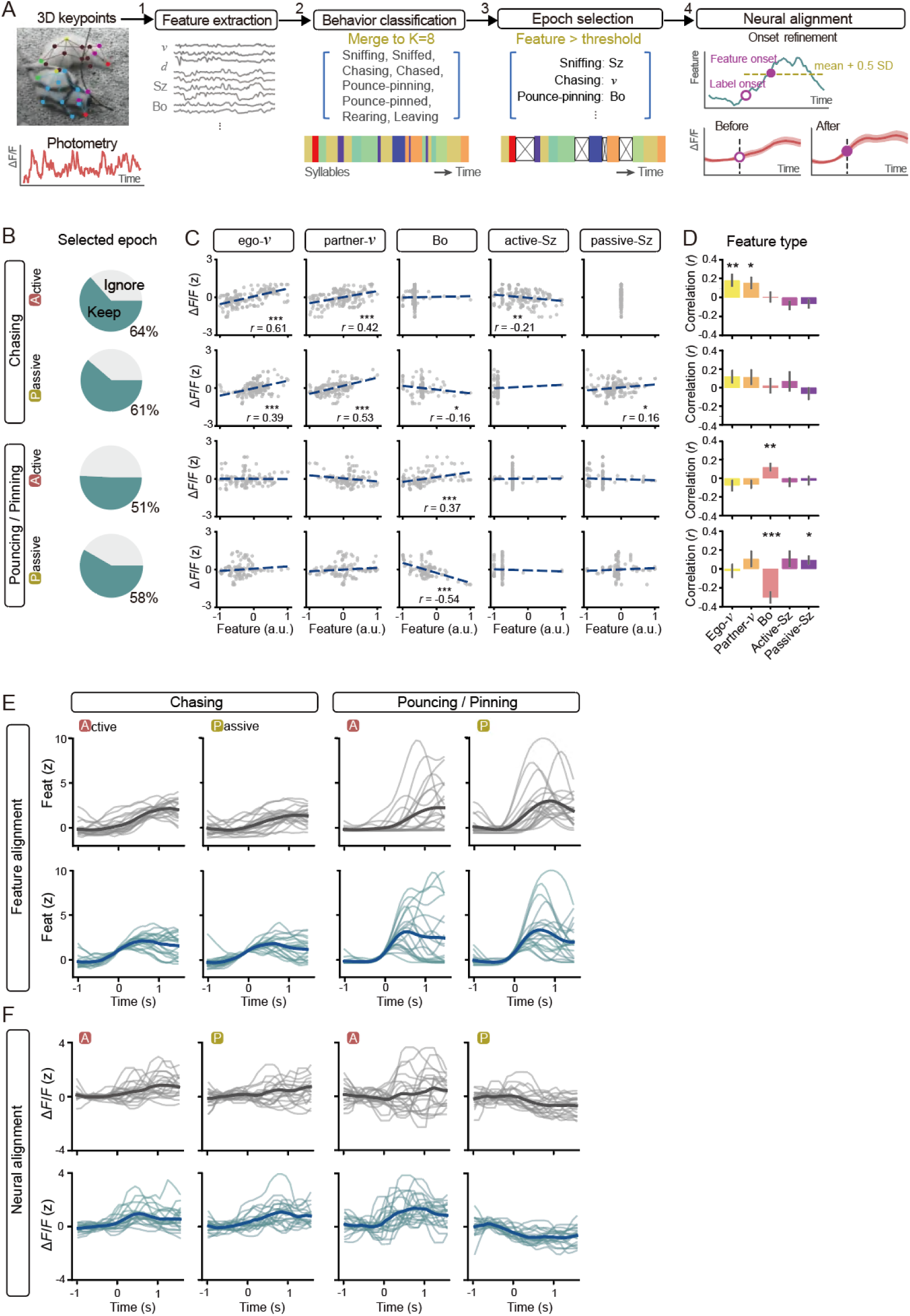
Behavior-specific dopamine dynamics and feature correlations, Related to Figure 4. (A) Behavior-dopamine alignment pipeline. (1) Feature extraction: Normalized behavioral features derived from 3D keypoint trajectories of interacting rat pairs (with rat implanted with an optical fiber). (2) Behavioral classification: Video clips were categorized into 36 behavioral classes using pretrained Seq2seq-FCN model and subsequently consolidated into 8 core behavioral modules (Table S2). (3) Epoch selection: Behavioral transitions with selected feature changes exceeding a predefined threshold (STAR Methods) were identified as representative epochs. Only sessions with≥3 representative epochs were included. (4) Neural alignment: DA signals were synchronized to feature onset (solid purple dot), defined as the first time point after behavioral label onset (open purple dot) when feature values exceeded 0.5 SD above the mean (dashed threshold line). (B) Percentage of selected behavioral epochs. Epochs with feature changes above a predefined threshold were included for (top) chasing dynamics (active *vs.* passive roles) and (bottom) play behaviors (active *vs.* passive roles). Only selected epochs were used in subsequent neural correlation analyses. (C and D) Feature-specific correlations with dopamine response dynamics. (C) Correlation coefficients (r values) between behavior-associated DA responses and key interaction features in a representative male rat dyad. (D) Correlations across multiple male pairs (n = 23). Analyzed features include: Ego-*v*: egocentric velocity; Partner-v: partner’s velocity; Bo: body occlusion area; Active-SZ: Active snout-to-passive tail region overlaps; Passive-SZ: Passive snout-to-active tail region overlaps. Data shown as mean ± SEM; statistical significance was assessed using two-sided paired t test (\**p* < 0.05, \*\**p* < 0.01, \*\*\**p* < 0.001). (E and F) Temporal dynamics of behavioral features and dopamine signals. (E) Fluctuations in behavioral features and (F) corresponding dopaminergic activity fluctuations across randomly selected representative behavioral epochs. Traces are aligned to behavioral label onset (top) and feature onset (bottom), respectively.

**Figure S5.**
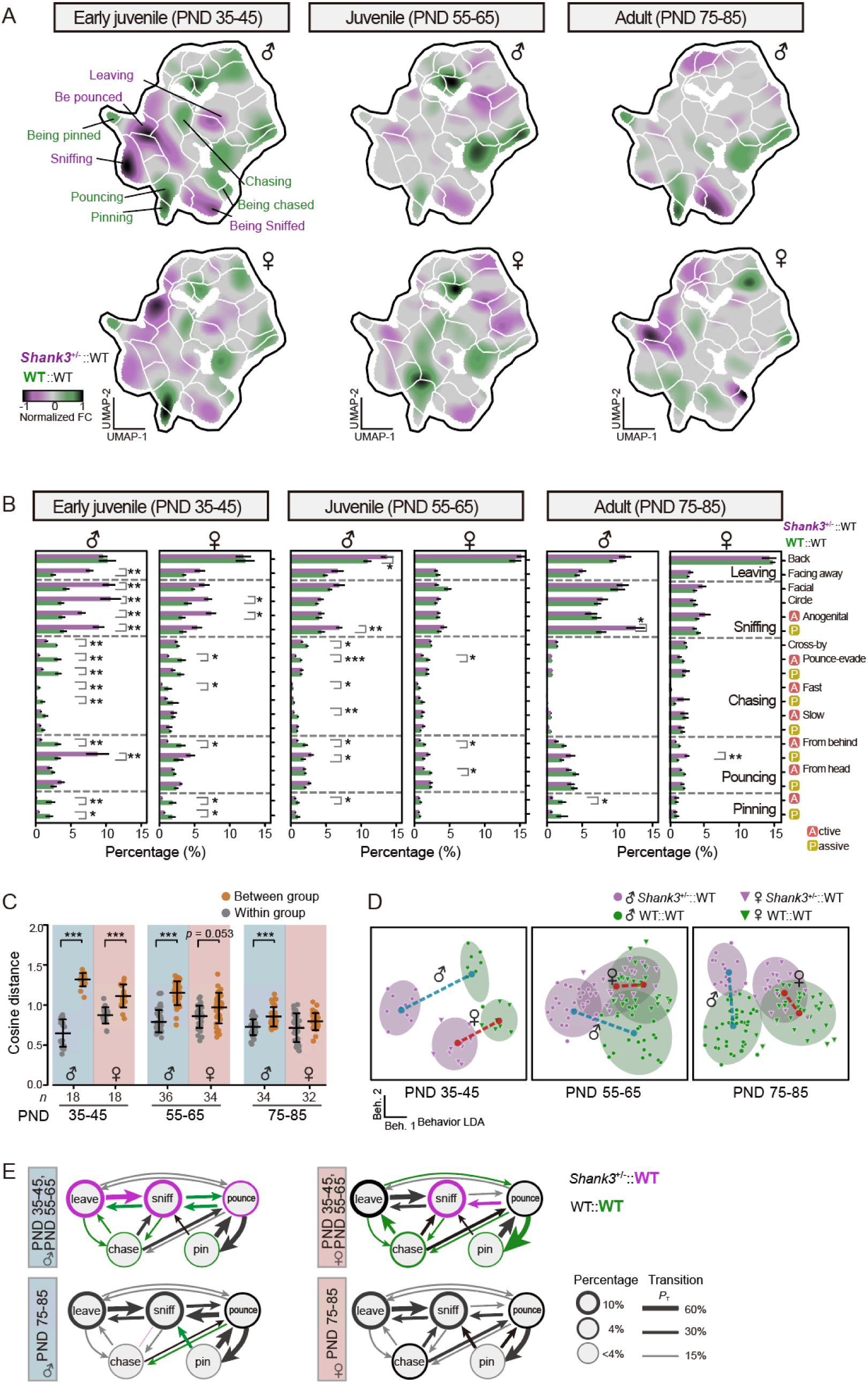
Behavioral characterization of *Shank3*^+/−^::WT rat pairs across development and sexes, Related to Figure 5. (A and B) Genotype comparison of behavioral clusters. (A) UMAP visualization of behavioral cluster occupancy differences between *Shank3*^+/−^::WT (purple) and WT::WT (green) dyads. (B) Proportional distribution of behavioral clusters (FDR-adjusted two-sided unpaired t test). (B) Behavioral profile similarity. Cosine distance analysis comparing intra-group (gray) versus inter-group (yellow) behavioral profiles (STAR Methods). Males: blue; females: red (FDR-adjusted two-sided paired t test). (C) Linear discriminant analysis (LDA) of behavioral profiles. Distinct clustering of *Shank3*^+/−^::WT (purple) and WT::WT (green) groups in males (circles) and females (triangles). Ellipses represent 90% confidence intervals. Dashed lines indicate pairwise distance between centroids, highlighting sex-dependent genotype discriminability (STAR Methods). (E) Behavior state transition diagrams comparing WT partners in *Shank3*^+/−^::WT pairs versus WT::WT pairs. Top/bottom: juvenile/adult; left/right: males/females. Significant differences (FDR-adjusted two-sided unpaired t test, *p* < 0.05) are color-coded. All data: mean ± SEM; \**p* < 0.05, \*\**p* < 0.01, \*\*\**p* < 0.001.

**Figure S6.**
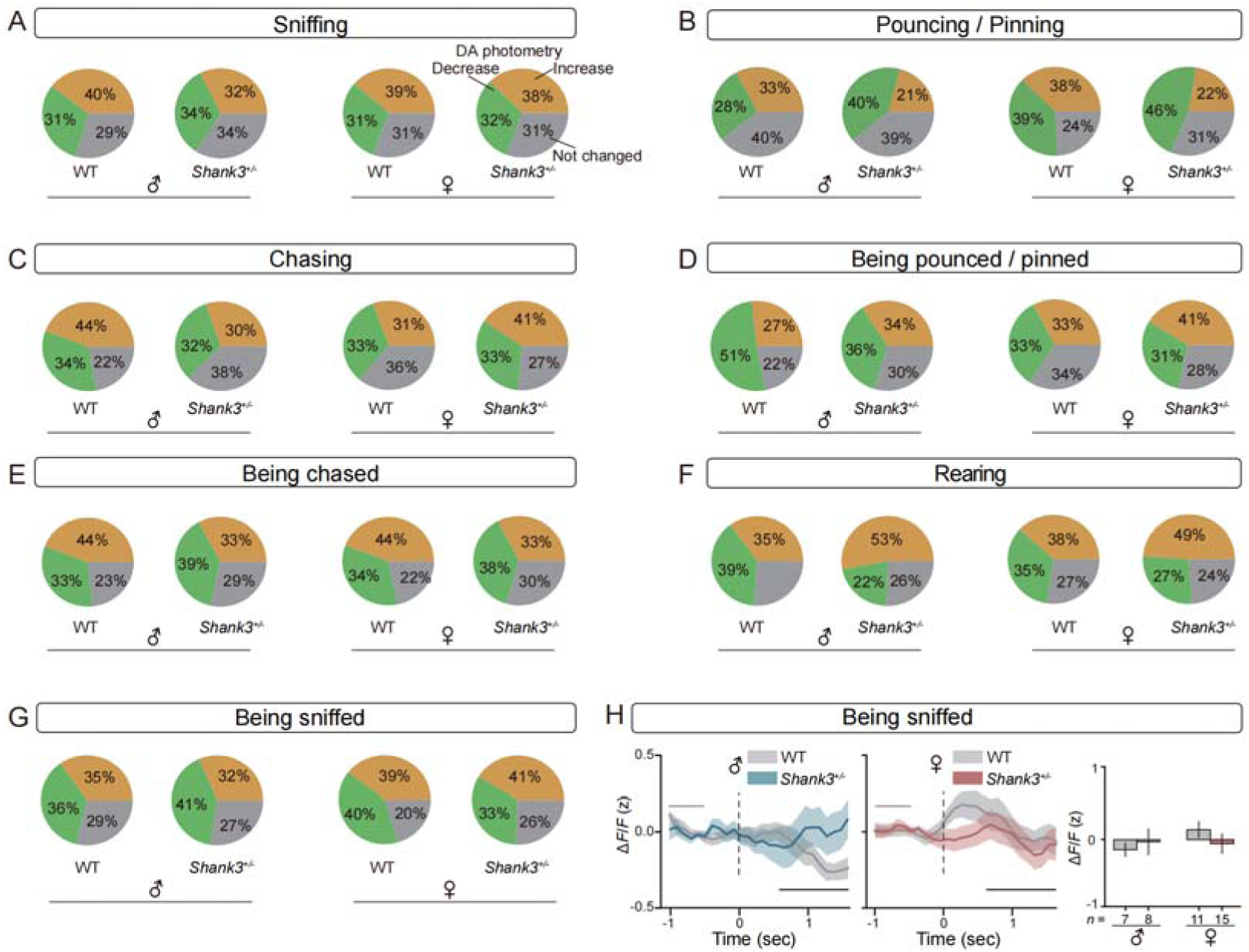
Dopamine response profiles during social interactions in WT and *Shank3*^+/−^juveniles, Related to Figure 6. (A-G) Proportions of behavioral epochs exhibiting significant DA increases (yellow), decreases (green), or non-significant changes (gray) in WT and *Shank3*^+/−^juveniles across different behavioral categories. The one-sided unpaired t test was conducted between baseline and response for each behavioral epoch using the same windows with Figure 6. (H) Time-resolved dopaminergic dynamics in passive sniffing. Left and middle (traces): Z-scored ΔF/F aligned to behavioral onset (t = 0; STAR Methods), shown as mean ± SEM (male: blue; female: red; ≥ 3 epochs/video). Gray line: baseline (−1 to 0.5 s). Black line: response window (0.5 to 1.5 s). Right (bar plot): Peak z-scored ΔF/F (response window *vs.* baseline).

**Figure S7.**
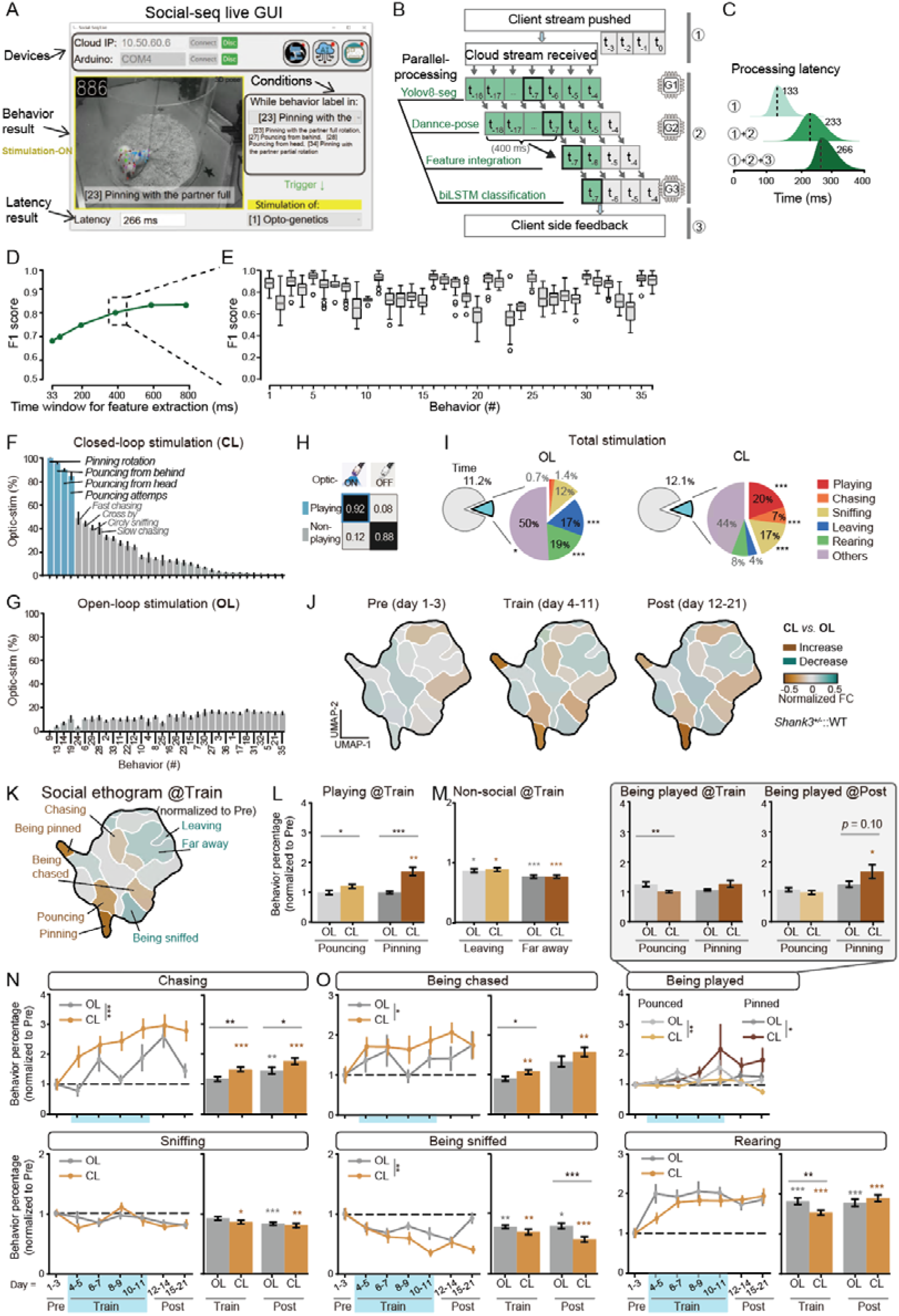
Closed-loop dopamine modulation during naturalistic social interactions, Related to Figure 7. (A) The graphical user interface (GUI) of the Social-seq platform. (B-C) Real-time processing pipeline. (1) Network transmission: Video frames captured at 120 fps were downsampled to 30 fps for cloud transfer. This initial transfer introduces introduced an averaged 4-frame delay (133 ± 12 ms) due to network latency. (2) Cloud computation: The cloud processing executes YOLOv8-based animal segmentation, DANNCE-powered 3D pose reconstruction, and BiLSTM behavior classification in parallel. This computational stage adds an averaged 3-frame delay (133 ± 16 ms). (3) Client-side feedback stage: Processed behavioral labels and corresponding video frames were returned to the client side with an additional average 1-frame delay (33 ± 8 ms). Latencies (mean ± SD) measured frame-by-frame in 15-minute. (D and E) Behavior classification performance. (D) F1 scores across different time windows used for real-time feature extraction. (E) The dashed box highlights the selected 400-ms window that provides an optimal balance between processing latency and classification accuracy across all 36 behavioral categories (n = 40 videos). (F and G) Optical stimulation distribution profiles. (F) Percentage of CL stimulation pulses allocated to each behavioral category. Blue bars highlight target play-related behaviors. (G) Distribution of OL stimulation across behavioral categories (n = 32 videos). (H) Proportion of stimulation specificity for target (blue) versus non-target (gray) behaviors in CL group. (I) Proportion of stimulation time allocated to different behavioral categories in CL and OL paradigms. (J) Cluster occupancy differences between CL and OL groups during pre-training, training and post-training phases (yellow: CL > OL; cyan: CL < OL). (K) Training-phase cluster occupancy normalized to pre-training baseline. (L and M) Duration of play-related (L) and non-social (M) behaviors during the training phase. (N and O) Behavioral dynamics. (N) Active social behaviors (chasing, sniffing) initiated by *Shank3*^+/−^juveniles. (O) Passive social behaviors and rearing. Line plots: Progressive behavioral changes (one-way ANOVA). Bar plots: Behavioral quantification in training and post-training phases (two-sided unpaired t test; CL: n = 32; OL: n = 30 videos). All data: mean ± SEM; \**p* < 0.05, \*\**p* < 0.01, \*\*\**p* < 0.001.

---

**Table S1.** Ethogram of Behavior Clusters.

| Cluster ID (#) |  | Interaction Type | Behavioral Description<br>(from the striped rat's perspective) |
| --- | --- | --- | --- |
| Refined<br>(K = 36) | Original<br>(K = 42) |  |  |
| 1 | 25 | Mutual | Back-to-back disengagement |
| 2 | 42 | Mutual | Mutual facial investigation |
| 3 | 11 | Mutual | Head-to-head approach |
| 4 | 31 | Mutual | Circular sniffing |
| 5 | 20 | Mutual | Parallel rearing (facing away) |
| 6 | 4 | Mutual | Head-to-head approaching |
| 7 | 32 | Mutual | Mutual rearing with facial investigation |
| 8 | 23 | Mutual | Back-to-back facing away |
| 9 | 10 | Reciprocal | Full rotation pinning |
| 10 | 28 | Reciprocal | Being fast chased |
| 11 | 37 | Reciprocal | Rearing while partner faces away |
| 12 | 17 | Reciprocal | Anogenital sniffing |
| 13 | 3 | Reciprocal | Posterior pouncing |
| 14 | 2 | Reciprocal | Frontal pouncing |
| 15 | 9 | Reciprocal | Being slowly chased |
| 16 | 35 | Reciprocal | Rearing facing partner's back |
| 17 | 18 | Reciprocal | Rearing during partner departure |
| 18 | 6 | Reciprocal | Posterior approaching |
| 19 | 15 | Reciprocal | Pouncing an evading partner |
| 20 | 5,7 | Reciprocal | Partial-rotation pinning |
| 21 | 16 | Reciprocal | Rearing while being observed |
| 22 | 30 | Reciprocal | Rearing within social proximity |
| 23 | 8 | Reciprocal | Being fully pinned |
| 24 | 13 | Reciprocal | Fast chasing |
| 25 | 19 | Reciprocal | Facing away from rearing partner |
| 26 | 38 | Reciprocal | Being anogenitally sniffed |
| 27 | 33 | Reciprocal | Being pounced from behind |
| 28 | 22 | Reciprocal | Being pounced head-on |
| 29 | 24 | Reciprocal | Slow chasing |
| 30 | 1 | Reciprocal | Rearing behind rearing partner |
| 31 | 12 | Reciprocal | Departing from rearing partner |
| 32 | 34 | Reciprocal | Being approached from behind |
| 33 | 40 | Reciprocal | Evading pounce attempts |
| 34 | 36 | Reciprocal | Being partially pinned |
| 35 | 14 | Reciprocal | Observing rearing partner |
| 36 | 14 | Reciprocal | Maintaining social range of rearing partner |
| - | 21,26,27,29,3<br>9,41 | Unreliable | Ambiguous rearing postures |

**Table S2.** Behavior consolidation and feature alignments.

| **Refined Categories**<br>(K = 36) | **Core Modules**<br>(K = 8) |  | **Kinematic Features for Onset Analysis** |
| --- | --- | --- | --- |
| 5, 7, 11, 16, 17, 21, 22, 30 | 1 | Rearing | Back height |
| 1, 31 | 2 | Leaving | Back-to-back distance |
| 26 | 3 | Being sniffed | Sniffing zone (target tail to partner nose) |
| 15, 10, 33 | 4 | Being chased | Partner's horizontal velocity |
| 23, 27, 28, 34 | 5 | Being pounce-pinned | Body occlusion area |
| 2, 3, 4, 12, 36 | 6 | Sniffing | Sniffing zone (target nose to partner tail) |
| 19, 24, 29 | 7 | Chasing | Egocentric horizontal velocity |
| 9, 13, 14, 20 | 8 | Pounce-pinning | Body occlusion area |
| 6, 8, 18, 25, 32, 35 | - | Others | - |

